# *S. cerevisiae* Cwc15p Tunes the Spliceosome Active Site for Remodeling and Catalysis

**DOI:** 10.64898/2026.03.20.713263

**Authors:** Natalie J. Zeps, Giovanni Balice, Elizabeth C. Baudhuin, Zach Freedman, Sierra Jones, Dennis Halterman, Aaron A. Hoskins

**Affiliations:** Department of Biochemistry, University of Wisconsin-Madison, Madison, WI 53706; Department of Plant Pathology, University of Wisconsin-Madison, Madison, WI 53706; USDA-ARS Vegetable Crops Research Unit, Madison, WI 53706; Department of Chemistry, University of Wisconsin-Madison, Madison, WI 53706

**Keywords:** pre-mRNA, splicing, yeast, spliceosome, RNA, *S*. *cerevisiae*

## Abstract

Pre-mRNA splicing is an essential step in eukaryotic gene expression during which spliceosomes remove introns from nascent RNAs while ligating the adjacent exons. Spliceosomes are cellular nanomachines composed of five small nuclear (snRNA) components and dozens of proteins, most of which are highly conserved. Despite the high conservation of many splicing factors between *S. cerevisiae* and *H. sapiens*, several protein components of the *S. cerevisiae* spliceosome are not essential for growth under normal laboratory conditions. This is particularly surprising for nonessential factors whose conserved domains contact the spliceosome’s catalytic core. Uncovering a function for these splicing factors can be challenging since they are not required for viability, may engage in functionally redundant interactions, and may display only weak phenotypes in the absence of secondary mutations in other spliceosome components. One such nonessential factor is the Cwc15 protein. Cwc15’s highly conserved N-terminus directly contacts the U2/U6 di-snRNA within the spliceosome catalytic core; yet its precise role in splicing has not been defined in any organism. In this work, we use molecular genetics in *S. cerevisiae* combined with splicing reporter assays to study Cwc15p function. We propose that Cwc15p not only promotes the stability of the active site but also its structural transitions. These functions may be critical for splicing in *S. cerevisiae* under nonoptimal conditions, facilitating use of weak or alternate splice sites, and could have implications for proofreading of spliceosome active site formation.

**Graphical Abstract:** 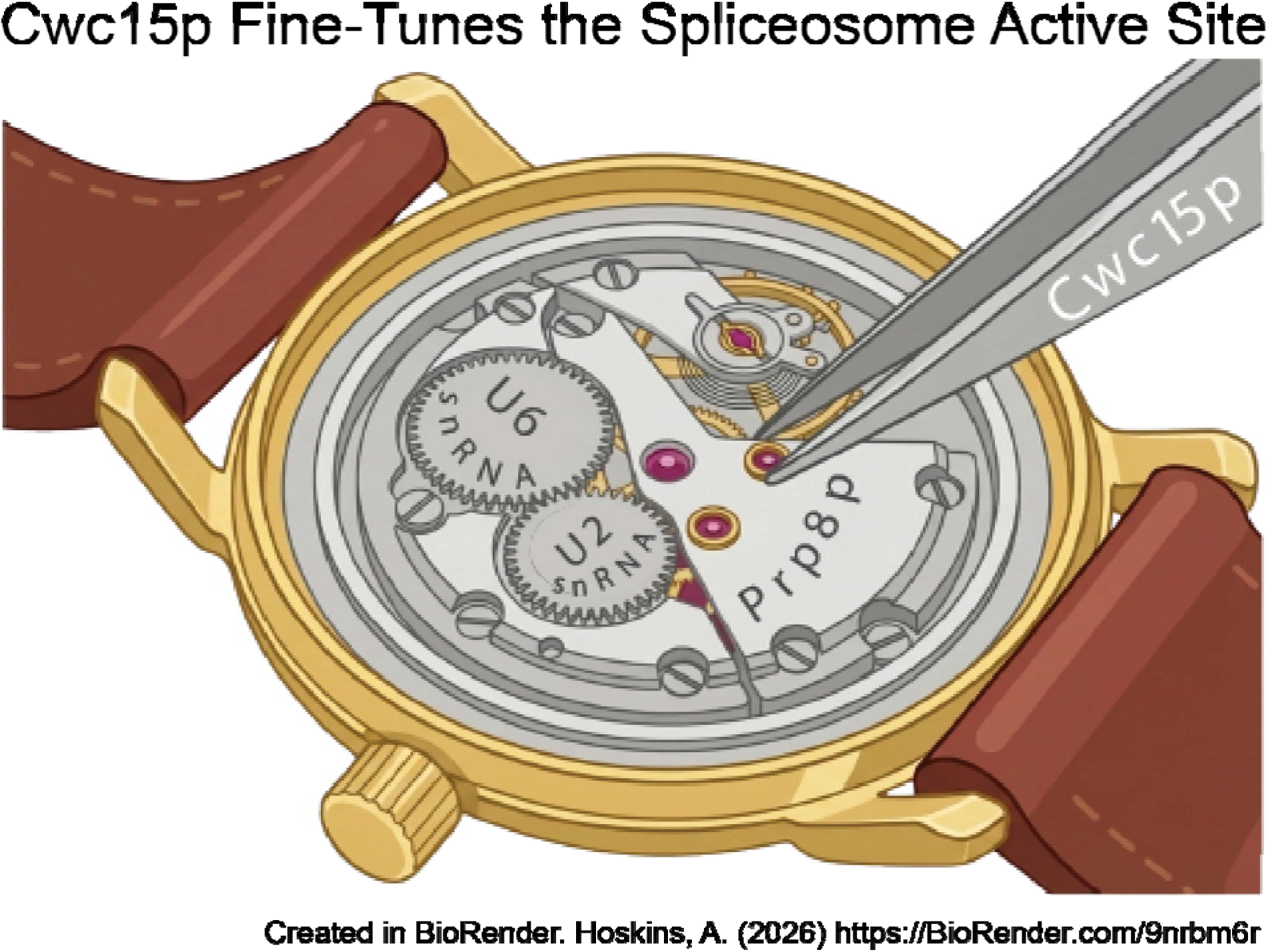

**Article Summary:** Pre-mRNA splicing is carried out by large macromolecular machines called spliceosomes which are composed of several snRNAs and dozens of proteins. Despite decades of study, the functions of many splicing factors such as *S. cerevisiae* Cwc15p remain unknown. Cwc15p is highly conserved among eukaryotes and directly contacts the spliceosome catalytic core. Here, we have used genetic and splicing reporter assays to study the function of Cwc15p during splicing *in vivo*. We propose that Cwc15p both stabilizes the spliceosome active site and impacts remodeling of that site.

## INTRODUCTION

In eukaryotes, nuclear DNA genes frequently contain introns which must be removed from precursor mRNA (pre-mRNA) transcripts concomitant with ligation of the flanking exons for gene expression (**Fig. 1A**). This process, called pre-mRNA splicing, is carried out by a large macromolecular machine called the spliceosome. Spliceosomes assemble around introns from uridine-rich small nuclear RNAs (the U snRNAs; U1, U2, U4, U5, and U6) and dozens of protein components. The snRNAs and many proteins associate with introns in pre-assembled complexes called small nuclear ribonucleoproteins (snRNPs). Other proteins also join spliceosomes in either multi-subunit complexes lacking RNA components [such as the Prp19-containing complex or Nineteen Complex (NTC)] or as individual units. The processes of spliceosome assembly and correct identification of introns are complex and involve numerous ATP-dependent and -independent stages (Plaschka et al. 2019; Kastner et al. 2019).

**Figure 1.**
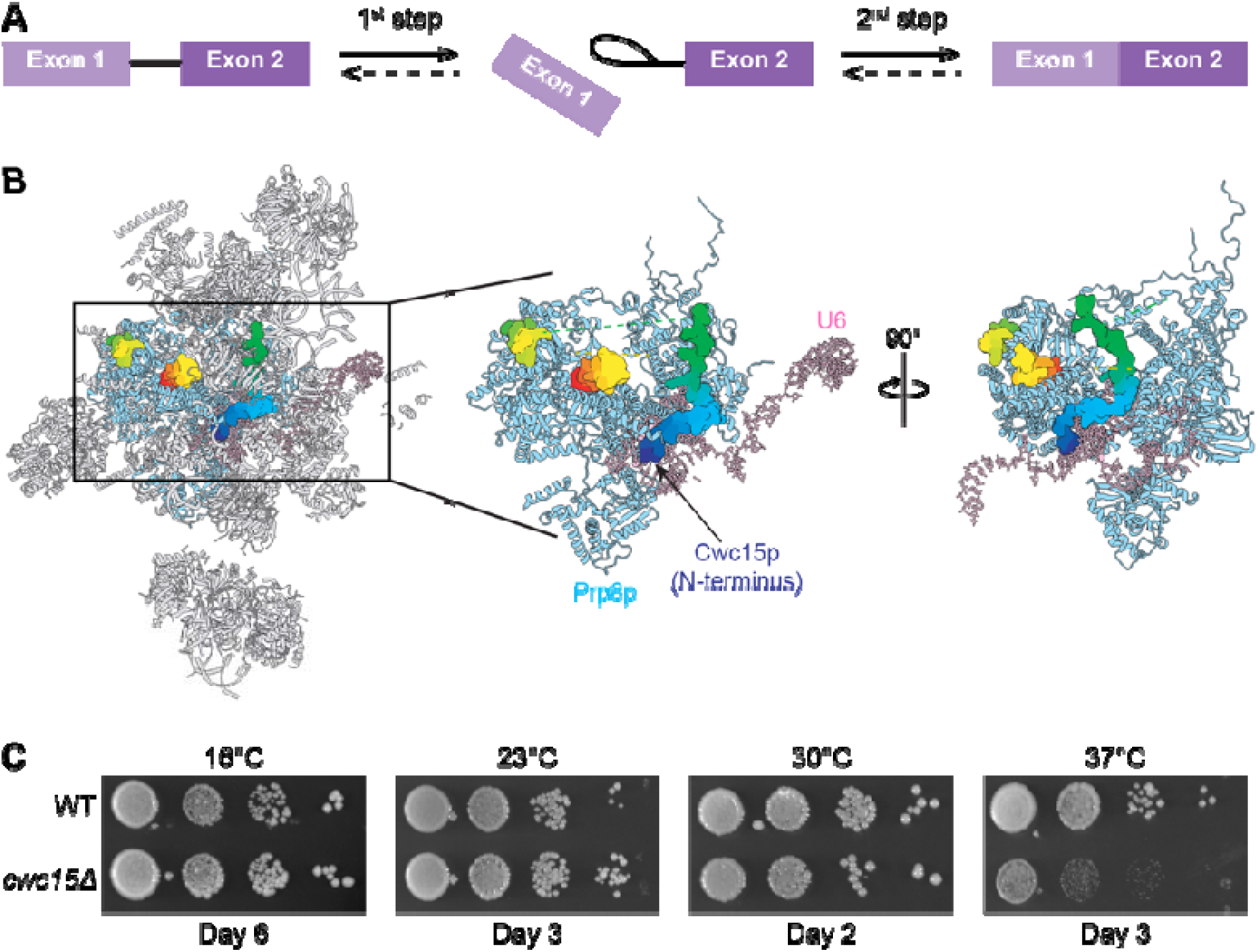
The transesterification steps in pre-mRNA splicing and location of Cwc15p in the spliceosome. (**A**) The two transesterification steps in splicing involve 5’SS cleavage (1^st^ step) and exon ligation (2^nd^ step). (**B**) Location of Cwc15p in the *S. cerevisiae* product (P) complex spliceosome (PDB 9DTR). This view shows the spliceosome in grey cartoon representation with Cwc15p shown in spacefill and colored with a gradient from blue (N-terminus) to red (C-terminus). Not all of the Cwc15p peptide chain could be modeled in the structure. Prp8p and U6 snRNA are also shown in blue and pink, respectively. A clipping mask was used to make the spliceosome active site and Cwc15p more visible. This figure was generated using ChimeraX (Meng et al. 2023). (**C**)Yeast without *CWC15* exhibit a *ts* phenotype at 37°C. For panel C, yeast were grown on YPD plates and imaged after the given number of days.

Once assembled, spliceosomes contain RNA active sites composed of the U2 and U6 snRNAs, the pre-mRNA substrate, and metal ions (Fica et al. 2013; Senn et al. 2024; Yan et al. 2019). Despite being ribozymes at their catalytic heart, the RNAs require numerous proteins to scaffold and arrange them into the catalytic conformations needed for 5’ splice site (5’SS) cleavage (1^st^ chemical step of splicing) and exon ligation (2^nd^ step). Some of these preoteins such as Prp8p, which scaffolds the snRNA active site (Grainger and Beggs 2005), are essential components of the splicing machinery. However, a number of other proteins are not essential (or at least required under certain conditions). In particular, the NTC and its related complexes contain many proteins that are non-essential in *Saccharomyces cerevisiae* (yeast) for growth and pre-mRNA splicing under laboratory conditions (Hogg et al. 2010). For several of these proteins, their non-essentiality is surprising given their locations near the spliceosome active site and contacts with critical protein and RNA components in cryo-EM structures. Network analysis of these structures suggested that non-essentiality may be due to redundancy in splicing factor interactions—non-essential factors make fewer unique interactions compared with essential factors (Kaur et al. 2022). This implies that they primarily function in *S. cerevisiae* to fine-tune splicing reactions under certain conditions or on certain substrates.

One nonessential component is yeast Cwc15p. Cwc15p is a small, 20 kDa protein that was identified as <u>c</u>omplexed <u>w</u>ith <u>C</u>ef1 (or cdc5 in *Schizosaccharomyces pombe*) by mass spectrometry of purified splicing factor complexes (Ohi et al. 2002). Despite being non-essential in *S. cerevisiae*, Cwc15p is highly conserved with homologs present throughout eukaryotes including in humans and plants (Ohi et al. 2002) (**Fig. S1**). Cwc15p homologs are essential in *S. pombe* (Ohi et al. 2002), *Arabidopsis* (Slane et al. 2020), and *Bos taurus* (Sonstegard et al. 2013). This suggests that organisms with more complex splicing regulatory mechanisms, intron composition, and/or machineries than *S. cerevisiae* require Cwc15p for proper spliceosome function (Sales-Lee et al. 2021).

No detailed biochemical or molecular genetics work has been reported for analysis of Cwc15p’s function during pre-mRNA splicing. Cryo-EM structures of catalytic-stage spliceosomes from a variety of organisms all show that Cwc15p lacks appreciable secondary structure and threads its way throughout the spliceosome (**Fig. 1B**). Cwc15p’s highly conserved N-terminus (**Fig. S1**) inserts directly into the catalytic core of the spliceosome, contacting the back of the catalytic U2/U6 snRNA duplex opposite of the face on which catalytically essential metal ions are bound (**Fig. 1B**, boxed region). Large portions of the remainder of Cwc15p are unresolved in available cryo-EM structures; however, a tryptophan in a conserved region near the C-terminus (Trp127, **Sup. Fig. S1**) is visible and interacts with a hydrophobic pocket on Prp8p. Given these interactions with critical components of the spliceosome, it is likely that Cwc15p influences pre-mRNA splicing despite being non-essential in *S. cerevisiae*. In support of this, *S. cerevisiae cwc15*Δ is synthetically lethal with the *prp19-1* allele (Ohi et al. 2002), and in *S. pombe*, the *fsm32* mutant allele of *CWF15* (the *S. pombe* homolog of *CWC15*) results in intron retention and splicing defects of the MCS2 transcript (Hoyos-Manchado et al. 2017). Additional evidence comes from studies of plant Cwc15 proteins. The *Arabidopsis cwc15-1* allele causes splicing defects *in vivo*, although these were not extensively characterized (Slane et al. 2020), and *S. tuberosum* (potato) Cwc15p was recently found to be the target of the pathogen *Phytophthora infestans* effector protein IPI-O1 (Sun et al. 2024). This interaction results in a change in splicing of an immunity gene in potato, resulting in pathogen resistance.

Here, we use yeast molecular genetics to probe the function of Cwc15p during pre-mRNA splicing. By testing genetic interactions between *cwc15*Δ and splicing factors and using *in vivo* splicing reporter assays, we propose that Cwc15p stabilizes the active site while also facilitating its remodeling.

## MATERIALS AND METHODS

### Yeast strain and plasmid construction

The *CWC15* gene was deleted from yeast strains by replacement with a hygromycin resistance cassette (HygR) through homologous recombination (Goldstein and McCusker 1999). Gene deletions were confirmed by PCR.

Mutant *CWC15* and *SNR6* plasmids were constructed either by site-directed mutagenesis using inverse PCR (Silva et al. 2023) or by purchasing plasmids containing the appropriate DNAs from GENEWIZ (South Plainfield, NJ USA) or Genscript (Pennington, NJ USA). All plasmids were sequenced before use by Sanger or whole-plasmid next-generation DNA sequencing (Functional Biosciences; Madison, WI USA). Yeast growth and transformation were carried out using standard protocols and media (Guthrie et al. 1991). Yeast strains and plasmids are listed in **Sup. Tables S1** and **S2**, respectively.

### Yeast growth assays at various temperatures

Yeast cultures were grown overnight in liquid culture to saturation and then adjusted to an OD_600_ of 0.5 in 10% (v/v) glycerol. Three consecutive 1:10 serial dilutions were performed by transferring 100 µL of culture into 900 µL of diluent, resulting in final dilutions of 1:1, 1:10, 1:100, and 1:1000. Diluted cultures were then spotted onto plates and incubated at 16°C, 23°C, 30°C, or 37°C before imaging with an ImageQuant™ LAS 4000 imaging system (GE Healthcare/Cytiva). Assays were performed in at least triplicate.

### ACT1-CUP1 assays

Yeast strains transformed with ACT1–CUP1 reporter plasmids were cultured in - Leu dropout (DO) media. Diluted cultures (OD_600_ = 0.5) were spotted onto -Leu DO plates supplemented with 0–2.5 mM CuSO_4_ (Lesser and Guthrie 1993; Carrocci et al. 2018) and incubated at 30°C for 3 days before imaging.

### Primer extension assays

Yeast were grown to mid-log phase in -Leu DO liquid media at 30°C and collected by centrifugation. Total yeast RNA was isolated using a MasterPure Yeast RNA Isolation Kit (Epicentre) following the kit directions except that two RNA pellets isolated from ∼1.0 OD_600_ of cell culture were resuspended in a total volume of 30 µL of RNase-free water (Ambion) and 0.5 µL Murine RNase Inhibitor (New England Biolabs) in order to achieve a high concentration of RNA. RNA (10 µg) was then added to a primer extension reaction containing primers complementary to the ACT1-CUP1 (5’/IRD700/GGCACTCATGACCTTC-3’, IDT) and U6 RNAs (5’/IRD700/GAACTGCTGATCATCTCTG-3’, IDT). Primer extension was carried out using Superscript III (ThermoFisher) and analyzed by denaturing polyacrylamide gel electrophoresis and fluorescence imaging as described (Kaur et al. 2020).

## RESULTS

### Genetic Interactions between *CWC15* and Spliceosomal ATPases

To assay Cwc15p function, we first deleted the *CWC15* gene from *S. cerevisiae* and studied yeast growth at various temperatures (**Fig. 1C**). Consistent with a recent report by Henke-Schulz and colleagues (Henke-Schulz et al. 2024), we observed that *cwc15*Δ yeast had no observable growth defect at 16, 23, or 30°C on YPD plates but grew more slowly and were heat sensitive (temperature sensitive or *ts*) at 37°C. (We note that the strength of the *ts* phenotype is condition and/or strain dependent, and yeast lacking *CWC15* can sometimes grow more robustly at 37°C, *c.f.* **Figs. 1C** vs. **Sup. Fig. 6**, for example).

Given Cwc15p’s location proximal to the spliceosome active site, we hypothesized that genetic interactions may exist between *cwc15*Δ and the DEAH-box ATPases responsible for remodeling the spliceosome during formation or disruption of the catalytic center. We tested genetic interactions between *cwc15*Δ and *PRP2*, *PRP16*, and *PRP22* (**Fig. 2A**). Prp2p is responsible for releasing U2 snRNP proteins from the branch site (BS)/U2 snRNA duplex and proofreading the catalytic core during the final stages of active site assembly (Schmitzova et al. 2023; Lardelli et al. 2010; Wlodaver and Staley 2014). Prp16p is responsible for releasing proteins from the spliceosome required for 5’SS cleavage in order to facilitate active site reorganization and for proofreading the 1^st^ step (Koodathingal et al. 2010; Chung et al. 2023; Tseng et al. 2011). Prp22p is responsible for proofreading exon ligation as well as releasing the mRNA product and proteins needed during the 2^nd^ step (Mayas et al. 2006; Company et al. 1991; Chung et al. 2025).

**Figure 2.**
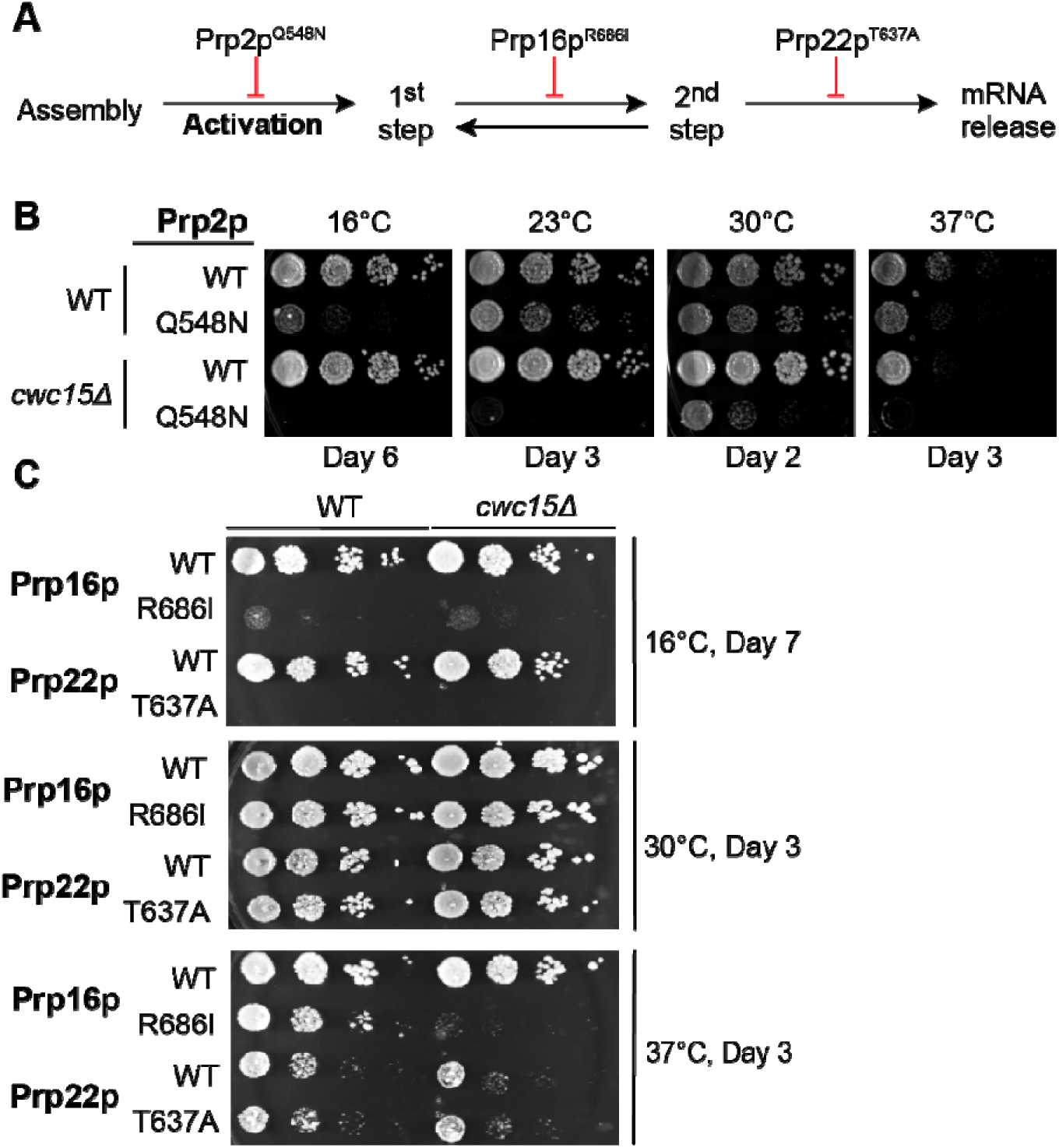
Genetic interactions between *CWC15* and spliceosome DEAH-box ATPases. (**A**) Schematic showing spliceosome remodeling events promoted by Prp2p, Prp16p, and Prp22p which promote activation, the 1^st^-to-2^nd^ step transition, and mRNA release, respectively. The mutations shown can inhibit these transitions and also impact pre-mRNA splicing fidelity. (**B**) Deletion of *CWC15* causes a negative genetic interaction with Prp2p^Q548N^. (**C**) Deletion of *CWC15* causes a negative genetic interaction with Prp16^R686I^ at 37°C. Yeast were grown on YPD plates and imaged after the number of days shown.

The Prp2p^Q548N^ mutation is cold sensitive (*cs*), inhibits splicing *in vivo*, and likely inhibits ATP binding or hydrolysis by the enzyme (Wlodaver and Staley 2014). Splicing inhibition arises due to difficulties in disruption of U2/U6 di-snRNA interactions needed for the final stages of catalytic site folding. Destabilizing mutations in these snRNAs or other splicing factors involved in activation can suppress the *cs* phenotype while stabilizing mutations can do the opposite (Wlodaver and Staley 2014; Carrocci et al. 2017; van der Feltz et al. 2021). When we combined Prp2p^Q548N^ with *cwc15*Δ, we observed that yeast were synthetic sick at 30°C and synthetic lethal at 16, 23, and 37°C (**Fig. 2B**). This indicates that Cwc15p functions during spliceosome active site formation and that loss of Cwc15p is detrimental if aberrant Prp2p activity is present. This phenotype could arise due to Cwc15p functioning to destabilize the U2/U6 di-snRNA during activation and/or stabilizing the product of Prp2p remodeling (*i.e.,* the 1^st^-step active site of the spliceosome).

We then tested genetic interactions with mutants in Prp16p and Prp22p. Prp16p^R686I^ is *cs* at 16°C and *ts* at 37°C, and at the latter temperature can be suppressed by the *prp8-156* allele which favors the exon ligation conformation of the active site (Query and Konarska 2004; Hotz and Schwer 1998). Similarly, the *cs* phenotype can be suppressed by *ecm2*Δ, which eliminates a nonessential splicing factor that makes stabilizing interactions in the 5’SS conformation (van der Feltz et al. 2021). Prp22p^T637A^ decouples ATP hydrolysis from RNA unwinding and causes a *cs* phenotype *in vivo* (Schwer and Meszaros 2000). Prp22p^T637A^ is also synthetic lethal at certain temperatures with loss of the 2^nd^-step splicing factor Fyv6p or alleles of *CEF1* that disrupt exon ligation (Lipinski et al. 2023; Query and Konarska 2012). Combined, these observations support a model in which Prp16p and Prp22p promote exit from the 1^st^ and 2^nd^-step conformations of the active site, respectively; Prp16p^R686I^ and Prp22p^T637A^ inhibit these transitions; and that the effects of these mutations can be suppressed (or exacerbated) by changes in the relative stabilities of the active site conformations (Liu et al. 2007).

Prp16p wild type (Prp16p^WT^)/*cwc15*Δ and Prp16p^R686I^/CWC15 yeast both grew at 37°C, but Prp16p^R686I^/*cwc15*Δ cells were very sick at this temperature and only tiny colonies were observed after 3 days (**Fig. 2C**, **Sup. Fig. S2**). In contrast, genetic interactions with Prp22p^T637A^ and *cwc15*Δ cells were not as apparent (**Fig. 2C, Sup. Fig. S2**). As with Prp2p, results with Prp16p are consistent with Cwc15p impacting spliceosome active site stability and in ways that change the ATPase-dependence of transitions between conformations or formation of transient, intermediate structures (Eysmont et al. 2019). Since Prp2p is involved in formation of the 1^st^-step active site and Prp16p is involved in its remodeling for the 2^nd^ step, these results suggest that Cwc15p is important for promoting transitions into and out of this spliceosome conformation.

### Genetic Interactions Between *CWC15, PRP8, and SNR6*

As mentioned above, different alleles or mutants of yeast Prp8p have been shown to influence pre-mRNA splicing by promoting the formation or stability of the active site for a given step at the expense of the other (*e.g.,* promoting the 1^st^ step reaction but inhibiting the 2^nd^ step, **Fig. 3A**) (Liu et al. 2007). Given our results with the ATPases (**Fig. 2**), we next tested genetic interactions between *PRP8* 1^st^- or 2^nd^-step alleles and *cwc15*Δ.

**Figure 3.**
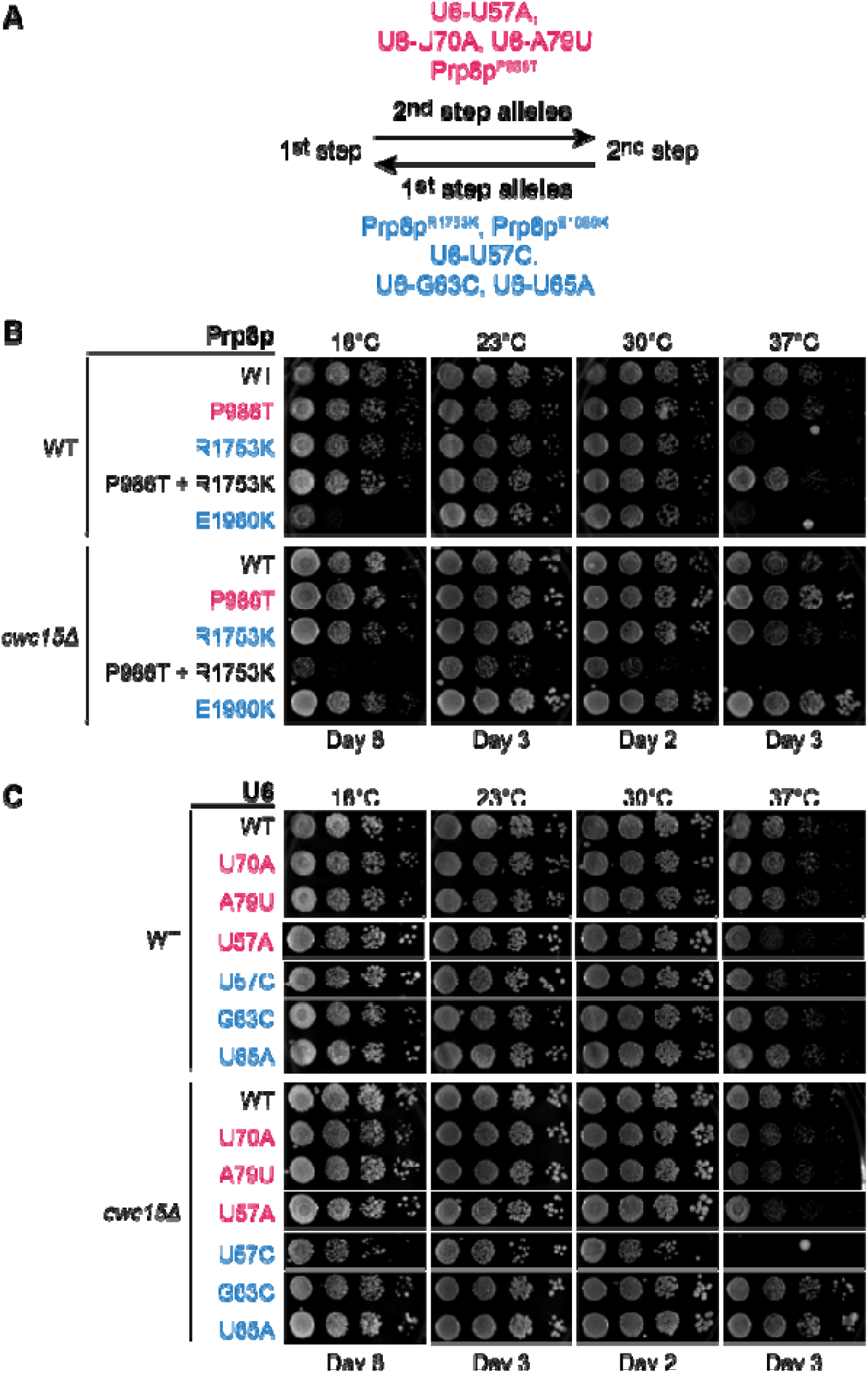
Genetic interactions between *CWC15* and *PRP8* or *SNR6* (U6 snRNA). (**A**) 1^st^-step alleles (blue) of Prp8p or U6 snRNA promote 5’SS cleavage and the 1^st^-step configuration of the spliceosome at the expense of the 2^nd^ step. 2^nd^-step alleles do the opposite (red). (**B**) Loss of *CWC15* can rescue the *ts* phenotype caused by Prp8 1^st^-step alleles. (**C**) Loss of *CWC15* is synthetic lethal with U6-U57C (1^st^-step allele) at 37°C and causes a *cs* phenotype at 16°C. For panels B and C, yeast were grown on YPD plates and imaged after the given number of days.

We expressed Prp8p mutants (Prp8p^P986T^, 2^nd^ step allele (*prp8-161*); Prp8p^R1753K^, 1^st^-step allele; and Prp8p^E1960K^, 1^st^-step allele (*prp8-101*)) (Liu et al. 2007) in the presence of *CWC15* or in a deletion background (**Fig. 3B**, **Sup. Fig. S3**). Deletion of *CWC15* was able to suppress the *ts* phenotype at 37°C for both 1^st^-step alleles (Prp8p^R1753K^ and Prp8p^E1960K^). Loss of *CWC15* was also able to suppress the *cs* phenotype of the Prp8p^E1960K^ mutant at 16°C. Any phenotypics differences in strains with Prp8p^P986T^ were not as apparent as *cwc15*Δ alone causes a *ts* phenotype even with Prp8p^WT^.

When 1^st^- and 2^nd^-step alleles are combined in the same *PRP8* gene, they can cancel each other’s effects (Liu et al. 2007). While the Prp8p^P986T+ R1753K^ double mutation grew at all temperatures in the presence of Cwc15p, yeast containing the double mutant Prp8 were both *cs* and *ts* when *CWC15* was deleted. These yeast grew either much more slowly (16°C) or not at all (37°C). This was unexpected since it indicates that the combination of the two mutations in Prp8p does not result in a protein with purely WT behavior with respect to Cwc15p.

The results with Prp8p^R1753K^ and Prp8p^E1960K^ are consistent with a model in which Cwc15p stabilizes the 1^st^ step conformation of the spliceosome active site at a stage that acts in concert with Prp8p 1^st^-step alleles. When the active site is destabilized due to loss of Cwc15p, the impact of *PRP8* 1^st^-step alleles is weakened and growth is restored at 37°C.

Since mutations in the U6 snRNA gene (*SNR6*) can also change the relative efficiencies of the 1^st^ and 2^nd^ steps splicing, we then tested whether U6 mutants that favor (U57A, U70A, A79U) or disfavor (U57C, G63C, U65A) the transition between the 1^st^ and 2^nd^ step also have genetic interactions with *cwc15*Δ (Liu et al. 2007; Query and Konarska 2004; Eysmont et al. 2019). These mutations also span different regions of the U6 snRNA including the upper and lower telestem (U70A, A79U and G63C, U65A; respectively) and the U2/U6 di-snRNA (U57A, U57C). The lower telestem is proposed to play a key role in formation of a transitional spliceosome structure between active conformations (Eysmont et al. 2019).

In this case, we saw the strongest genetic interactions between U6-U57C and *cwc15*Δ, which were synthetic lethal at 37°C and caused slow growth at 16°C relative to cells with *CWC15* (**Fig. 3C**). U57C is thought to stabilize the 1^st^ step conformation of the spliceosome or disfavor transitions away from this state (Liu et al. 2007; Eysmont et al. 2019; Wlodaver and Staley 2014). We favor an interpretation that Cwc15p facilitates the 1^st^-to-2^nd^ step transition and loss of Cwc15p in combination with U6-U57C makes this transition more difficult. In addition, milder negative growth phenotypes at 37°C were also observed for the U6-G63C, U70A, and A79U mutants in the presence of *cwc15*Δ (**Sup. Fig. S4**). In these cases, yeast lacking *CWC15* grew more slowly than those with the gene, and this was most apparent after 2 days of incubation at 37°C (**Sup. Fig. S4**). It is possible Cwc15p may generally help to stabilize the spliceosome active site when U6 snRNA mutations are present (also see **Fig. 6B**).

In combination with results from Prp8p and the ATPases, one hypothesis could be that Cwc15p both stabilizes the active site during 5’SS cleavage and facilitates rearrangement of the active site at stages preceding or following catalysis. The latter activity could occur by destabilization of the active site conformation at those times and/or by impacting the formation or stability of structures that form during remodeling. Importantly, these data do not exclude indirect effects of Cwc15p on the active site in which the protein may also influence the equilibria (or kinetic stabilities) between different active site conformations rather than imparting structural stability directly. The negative genetic interactions between *cwc15*Δ and Prp8^P986T+R1753K^ also suggest that interactions between Cwc15p and Prp8p may be complex and potentially impact multiple steps and transitions in different ways.

### Impact of *CWC15* Deletion on Yeast Growth with ACT1-CUP1 Splicing Reporters

To further assess Cwc15p’s role in pre-mRNA splicing, we used the *in vivo* ACT1-CUP1 splicing reporter assay (Lesser and Guthrie 1993). In this assay, copper-sensitive yeast are transformed with a plasmid encoding a gene for the Cup1p copper resistance protein with derivatives of the *ACT1* intron inserted into its coding sequence (**Fig. 4A**). They are then grown on plates containing increasing concentrations of Cu^2+^. Growth in the presence of copper requires splicing of the *ACT1-CUP1* pre-mRNA and expression of the Cup1p protein. The degree of resistance of the yeast to copper correlates with the extent of splicing.

**Figure 4.**
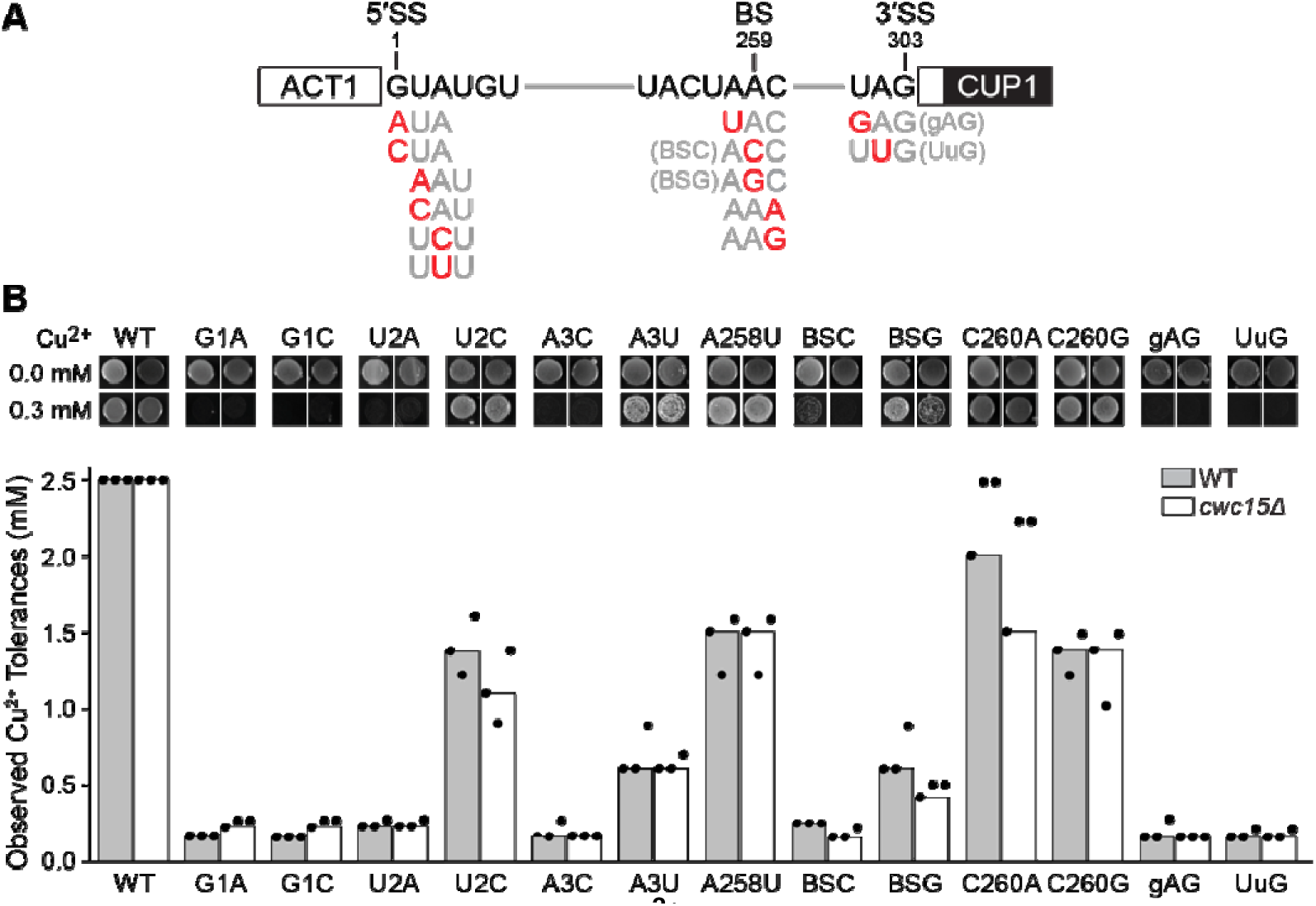
Loss of Cwc15p impacts Cu^2+^ tolerances of yeast expressing non-consensus ACT1-CUP1 reporters. (**A**) Schematic of the ACT1-CUP1 reporter pre-mRNA with the tested substitutions shown. (**B**) Loss of Cwc15p changes copper tolerances for yeast expressing reporter pre-mRNAs with particular substitutions within the 5’SS or BS but not for those expressing a reporter with consensus sites (WT). Bar heights represent values obtained from a single replicate of the ACT1-CUP1 assay. Dots represent values obtained for all three replicates. Yeast were grown at 30°C on - Leu DO plates containing various [Cu^2+^] and imaged after 3 days.

Using the WT ACT1-CUP1 reporter, which contains consensus yeast splice sites, we observed no change in resistance to copper between WT and *cwc15*Δ strains and both readily grew on plates containing the maximal copper concentration (2.5 mM) (**Fig. 4B**). This indicates a high splicing efficiency for the ACT1-CUP1 reporter even when Cwc15p is absent under these conditions. This likely explains, at least in part, the lack of a significant growth phenotype at 30°C for the deletion strain since the majority of yeast introns contain consensus splice sites.

We next tested growth of yeast containing reporters with a variety of substitutions at the 5’ or 3’SS or BS sequence (**Fig. 4A**, **B**). In these cases, we saw changes in copper tolerances for the WT *vs. cwc15*Δ strains for particular variants. For the 5’SS, we observed slight increases in tolerance for G+1 substitutions (G1A, G1C) and a reduction in tolerance for the U2C reporter. For substitutions at the branch site, we observed reductions in tolerance for both the BS-C and BS-G substitutions as well as for a C260A substitution at the 3’ flanking position. We did not observe any changes in copper tolerance for yeast with substitutions at the −3 (gAG) and −2 (UuG) positions of the 3’SS.

Primer extension analysis showed similar levels of RNA and splicing efficiencies for both strains with the WT reporter and lower levels of mRNA being produced with the U2C and BS-G reporters in the *cwc15*Δ strains (**Sup. Fig. S5**). However, the primer extension assay was not sensitive enough to confirm changes in mRNA production for the G1A or G1C reporters, and the assay showed similar levels of mRNA being produced with the C260A reporter.

Overall, the results show that loss of *CWC15* can impact the splicing reaction and lead to changes in both copper tolerance and mRNA production for some reporters. Despite impacting growth due to the variant ACT1-CUP1 reporters, the *cwc15*Δ allele does not behave identically to characterized Prp8 1^st^ or 2^nd^-step alleles since we observe small changes at the G+1 position but not other characteristic changes in growth (*e.g.*, decreased copper tolerance for the A3C reporter for 1^st^-step alleles and increases in copper tolerances for gAG and UuG reporters for 2^nd^-step alleles) (Liu et al. 2007; Query and Konarska 2004).

### Genes Encoding Mutant Cwc15 Proteins Can Rescue Yeast Growth

We next tested the importance of conserved regions of Cwc15p. The most highly conserved region in Cwc15p is the N-terminus, which inserts into the spliceosome active site between the U2 and U6 snRNAs and contacts many key splicing factors (**Fig. 5A**, **B**). In a cryo-EM structure of the *S. cerevisiae* spliced product spliceosome (P complex), Cwc15p residues Thr2 and Thr3 form hydrogen bonding interactions with U6 snRNA nucleotides C84 and C85 (Senn et al. 2024). Arg6 also forms a cation-π stacking interaction with U2 snRNA G20, and Gly14 closely packs into an interface with Prp46p. Additionally, conservation of Pro7 may be due to limiting flexibility of the N-terminus and facilitating its threading through a small channel formed between U2 snRNA G20/G21 and U6 snRNA A62/G63. We hypothesized that these interactions could be disrupted using T2V/T3V substitutions which replace hydrophilic with hydrophobic side chains that cannot hydrogen bond to U6 nucleotides, an H5A/R6A/P7A/Q8A quadruple mutant (AAAA) that eliminates the cation-π interaction with U2 and increases flexibility at position 7, or by introducing a side chain at the interface with Prp46p with a G14A mutation. Finally, we also deleted the first thirteen amino acids of Cwc15p entirely (ΔNterm).

**Figure 5.**
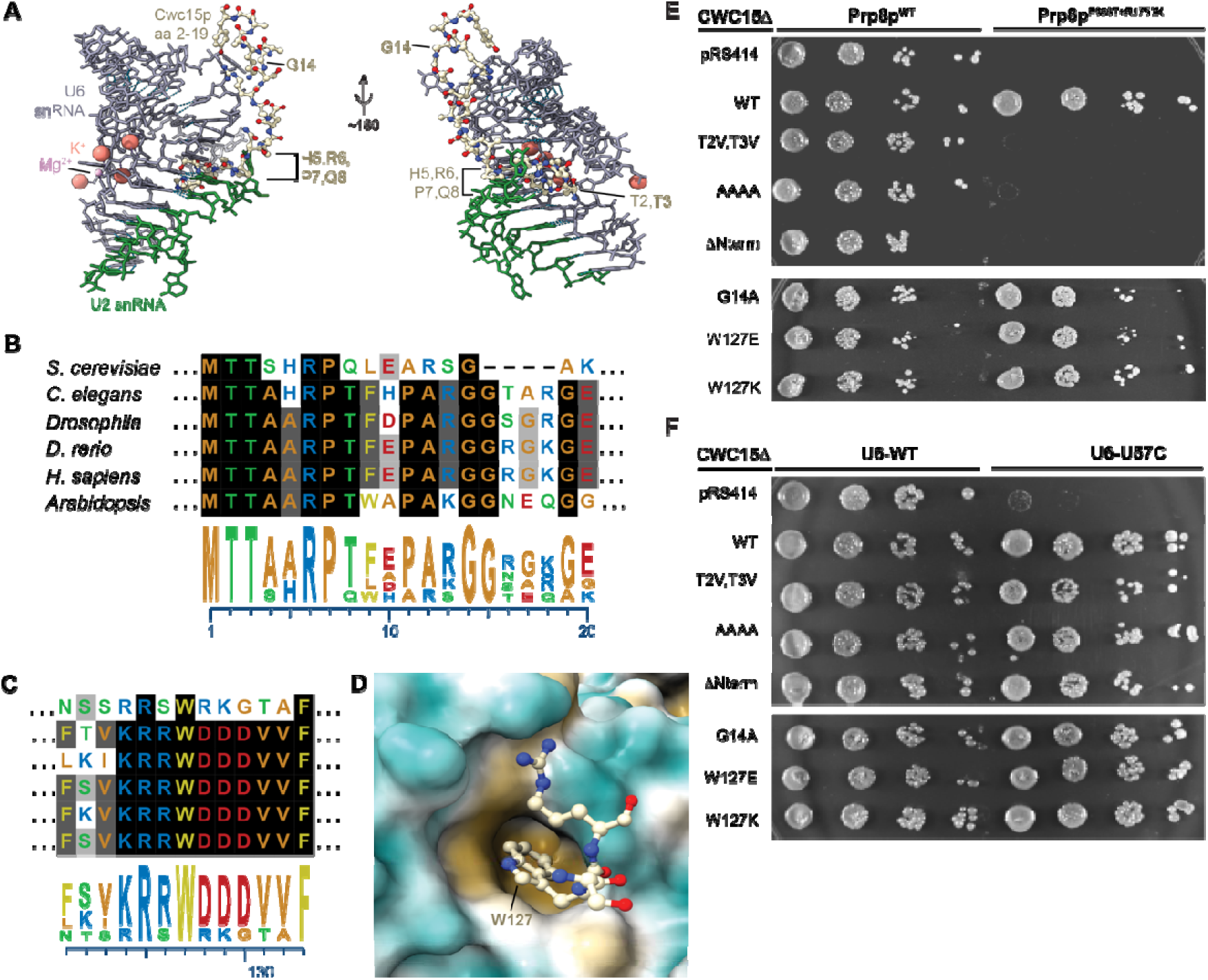
Cwc15p mutants can complement a *cwc15*Δ growth defect. (**A**) Close-up of Cwc15p N-terminus interactions with the U2/U6 di-snRNA within the active site of a P complex spliceosome (PDB 9DTR). The active site can be identified by the presence of Mg^2+^ and K^+^ ions. (**B**) Sequence alignments and logos for Cwc15 from single and multicellular model organisms and humans. The N-terminus of Cwc15 is highly conserved among eukaryotes. (**C**) Trp127 is also highly conserved. (**D**) Close-up of the interaction of Cwc15p residue Trp127 with a hydrophobic pocket on Prp8p (surface representation, hydrophobic surfaces shown in yellow and hydrophilic in blue-green). (**E, F**) Growth assays of *cwc15*Δ yeast expressing Cwc15p mutants relative to WT and empty plasmid (pRS414) controls in the presence of Prp8p or U6 mutants. Yeast were grown on -TRP DO plates and imaged after 2 days. Panels A and D were created using ChimeraX (Meng et al. 2023). Sequence alignments and logos for panels B and C were generated using MegAlign Pro (Lasergene, DNASTAR, Inc.).

In addition to the N-terminus, Cwc15 proteins also contain a conserved tryptophan residue located in the C-terminal half (**Sup. Fig. S1**, Trp127). This tryptophan is flanked by charged amino acids and fits into a hydrophobic pocket found on the Prp8 protein in the reverse transcriptase-like domain (**Fig. 5C, D**). Therefore, we also made Cwc15p mutants in which the tryptophan was substituted with a differently-shaped and charged residue (glutamate or lysine), which would be less favorable to bury within this pocket.

We transformed a *cwc15*Δ yeast strain with TRP1/CEN plasmids containing either WT Cwc15p or mutants expressed from the endogenous promoter and selected for growth on -trp plates. None of the mutants caused any growth defect at 30°C (not shown). Suprisingly, all of the mutants, included the N-terminal deletion, were able to rescue the *ts* growth phenotype of *cwc15*Δ yeast at 39°C (**Fig. S6**).

We then tested if the mutants could rescue lethal phenotypes observed at 37°C when *cwc15*Δ was combined with either Prp8p^P986T+R1753K^ or U6-U57C alleles (**Fig. 5E**, **F**). In the case of Prp8p^P986T+R1753K^, we observed mutant-specific effects. While all of the Cwc15p mutants rescued growth of U6-U57C yeast at 37°C, only the G14A, W127E, and W127K mutants rescued growth with Prp8p^P986T+R1753K^. Thus, growth in the presence of Prp8p^P986T+R1753K^ is dependent on the conserved N-terminus of Cwc15p but not on Gly14 or Trp127. Rescue of U6-U57C is not dependent on the Cwc15p N-terminus even though these amino acids contact the U6 snRNA in the active site of the spliceosome.

Rescue of the deletion or U6-U57C *ts* phenotypes could depend on spliceosomal interactions not yet structurally defined or Cwc15p’s interactions with other machineries such as the transcription and export (TREX) complex (Henke-Schulz et al. 2024). Cwc15 has not been fully resolved in any spliceosome structure from any organism— the structural environment of large portions of the protein are unknown. For example, in the most highly resolved spliceosome structure from *S. cerevisiae*, 58% of the Cwc15p peptide could not be modeled (Senn et al. 2024).

### Genetic Interaction Between *CWC15* and Destabilizing U6 snRNA Mutants

Finally, we further tested the idea that Cwc15p stabilizes the active site by introducing destabilizing mutations into the U6 snRNA in regions that contact the N-terminus of Cwc15p (**Fig. 6A**). We introduced U6 A62U, G63U, C84G, or C85G mutations since it was previously shown that these U6 variants have little effect on yeast growth by themselves (McPheeters 1996). When the growth of *CWC15* and *cwc15Δ* yeast were compared with these variants, we observed a synthetic sick interaction *(i.e*., slower growth) between *cwc15*Δ and the U6 snRNA A62U mutation at 37°C. It is not obvious what structural explanation could account for this. One hypothesis is that Cwc15p could prevent inappropriate wobble pairing between U6 A62U and U2 nucleotides G20 or G21 in precatalytic or catalytic spliceosomes (Senn et al. 2024; Galej et al. 2016; Rauhut et al. 2016). The conformation of U2 G20 may be particularly important for proper active site formation (see **Sup. Fig. S7** and Discussion). For the remaining U6 snRNA mutations studied here, it is likely that they were not as significantly destabilizing and lack observable Cwc15p-dependent phenotypes in these assays.

**Figure 6.**
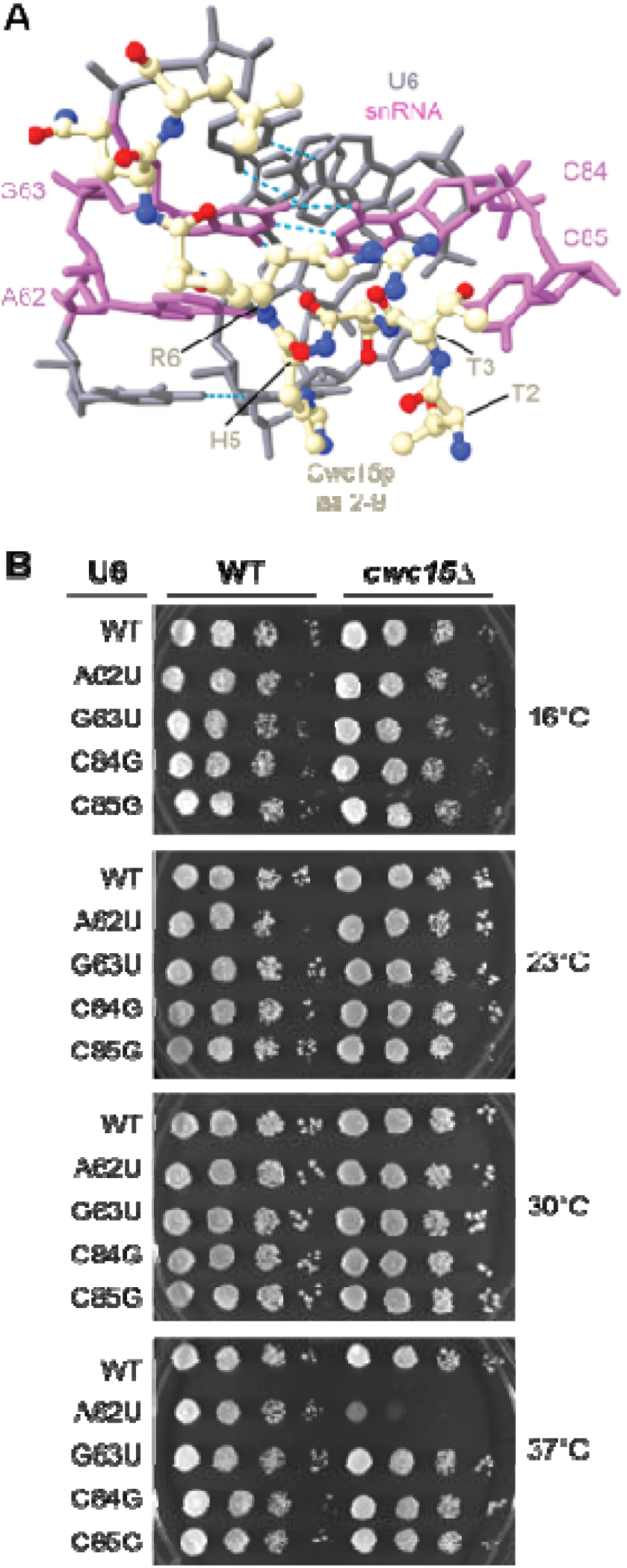
Genetic interactions between potentially active site-destabilizing mutations in the U6 snRNA and *CWC15*. A negative genetic interaction is observed between U6 snRNA A62U and *cwc15*Δ at 37oC. Yeast were grown on YPD plates and imaged after 2 (30°C), 3 (23 and 37°C), or 6 days (16°C). Panel A was created using ChimeraX (Meng et al. 2023).

## DISCUSSION

In summary, our data support a model in which Cwc15p fine-tunes the spliceosome active site and also impacts transitions between spliceosome conformations. The notion that non-essential *S. cerevisiae* splicing factors like Cwc15p perturb catalytic conformations of the spliceosome or enforce fidelity has previously been discussed with respect to other components of the NTC and the Ecm2 and Fyv6 proteins (Hogg et al. 2010; Senn et al. 2024; Lipinski et al. 2023; van der Feltz et al. 2021). Our group has previously proposed that non-essentiality may be linked to the relatively low number of unique connections proteins like Cwc15p make between essential factors and built-in interaction redundancy in the spliceosome (Kaur et al. 2022). Despite this, many non-essential splicing factors become critical at low or high temperatures or in certain genetic backgrounds. This could indicate that their conservation is due to subtle modulation of the splicing machinery or the need for additional structural stability under non-optimal growth conditions.

Beyond *S. cerevisiae*, this modulation may be critical for regulating alternative splicing and result in Cwc15 essentiality. One aspect of this model is potential adaptation (or exploitation) of Cwc15 for changing splicing outcomes in response to cellular conditions. For example, direct interaction between the *P. infestans* virulence protein IPI-O1 and the potato host protein StCwc15 leads to relocalization of StCwc15 from the nucleoplasm to the nucleolus and nuclear speckles (Sun et al. 2024). This interaction also results in a change in splicing of the *RB* immunity gene pre-mRNA leading to a decrease in the intron-retained isoform and an increase in the fully-spliced mRNA (Sun et al. 2024). While the molecular mechanism for this switch has not yet been determined, other NTC components are also subject to dynamic regulation including by post-translational modification (Lu and Legerski 2007; Lleres et al. 2010; de Almeida and O’Keefe 2015). This provides a plausible mechanism by which external factors could modulate Cwc15’s activity and, consequently, splicing efficiency.

Determining a more precise role for Cwc15p during splicing is challenging, and we hypothesize that the data are consistent with the protein having at least two functions: stabilizing the 1^st^-step active site during catalysis and promoting active site structural transitions. Yeast with Cwc15p show increased tolerance to Cu^2+^ using ACT1-CUP1 splicing reporters harboring a 5’SS U2C substitution and several BS substitutions (**Fig. 4, Sup. Fig. S5**). Additionally, loss of Cwc15p suppresses growth defects associated with Prp8p 1^st^-step alleles (**Fig. 3**) and results in a growth defect in the presence of a likely destabilizing U6 snRNA mutation (**Fig. 6**). These data not only suggest that Cwc15p could stabilize the 1^st^-step conformation of the spliceosome but that this may also be particularly important when non-consensus substrates or mutant splicing factors are present. An important caveat to this hypothesis is that it is difficult from genetic assays alone to disentangle direct active site stabilization/destabilization with indirect effects that perturb spliceosome dynamics or conformational equilibria during the reaction.

Among our results, genetic interactions between *cwc15*Δ, Prp2p^Q5548N^, Prp16p^R686I^, and U6-U57C mutations are particularly interesting. *CWC15* deletion has a negative genetic interaction with each of these mutations. While interpretation of U6 U57C mutations can be complex (Wlodaver and Staley 2014; McPheeters 1996; Query and Konarska 2004; Liu et al. 2007; Eysmont et al. 2019), the data are consistent with Cwc15p also facilitating structural transitions into and out of the 1^st^-step active site conformation. How exactly this occurs is unclear but could involve directly destabilizing the RNA structures that form the spliceosome active site, stabilizing transient active site conformations, or favoring formation of the stable products of these transitions. The latter possibility would provide a parsimonious explanation for some of our results: stabilization of the 1^st^-step active site by Cwc15p facilitates both its formation/remodeling by ATPases and 5’SS cleavage itself. At this time, we cannot yet conclude if similar activities for Cwc15p may also be present during the 2^nd^ step or other stages in splicing.

An intriguing possibility is that Cwc15’s conserved N-terminus participates in the above processes directly. Analysis of multiple cryo-EM structures of *S. cerevisiae* spliceosomes reveals that the very N-terminus of Cwc15p is located in between the U2 and U6 snRNAs in the B* spliceosome (the stage immediately preceding 5’SS cleavage) and in this same position in subsequently formed complexes (B* C C* P ILS complex spliceosomes) (Senn et al. 2024; Wan et al. 2017; Fica et al. 2017; Wan et al. 2019; Wilkinson et al. 2021) (**Sup. Fig. S7**). Importantly, in activated spliceosomes (B^act^) trapped prior to Prp2p activity and formation of B*, this same region of Cwc15p could not be modeled between the snRNAs and does not adopt a specific, stable conformation (see **Sup. Fig. S7**) (Bai et al. 2021). Further, the N-terminus of Cwc15 proteins could not be modeled in structures of an off-pathway, B^act^-like *S. cerevisiae* spliceosome assembled on an excised intron (Li et al. 2026) (Max Wilkinson, personal communication); a human spliceosome trapped while Prp2 was functioning (Schmitzova et al. 2023); or in *S. pombe* and *Chaetomium thermophilum* spliceosomes trapped prior to the B* stage and in the midst of being disrupted by the DHX35 ATPase (Soni et al. 2025; Li et al. 2025). Stable insertion of the Cwc15 N-terminus in between the U2 and U6 snRNAs may be a consequence of Prp2 activity and a marker of a correctly assembled active site. Conversely, a dynamic Cwc15 N-terminus (or just lack of its insertion between U2 and U6) may facilitate active site disruption and proofreading by DHX35. These putative functions are evidently not essential in *S. cerevisiae*, in which a corresponding DHX35-like activity has also not yet been identified.

The involvement of Cwc15p in other steps in gene expression besides pre-mRNA splicing complicates phenotypic analysis in our assays. In addition to splicing, Cwc15p is also important for tethering the TREX complex to the NTC, which itself is involved in both pre-mRNA splicing and transcription elongation (Henke-Schulz et al. 2024; Chanarat et al. 2011). Further, loss of *CWC15* has a negative genetic interaction with deletion of the general transcription factor TFIIS (*dst1*Δ) (Henke-Schulz et al. 2024). Given the high degree of coupling between transcription and splicing (Carrocci and Neugebauer 2024), it is likely that the growth phenotypes reported here have contributions from both processes. Future deconvolution of Cwc15p function may benefit from both molecular and genetic approaches to disentangle these processes as well as studies in model organisms in which the protein is essential for viability.

## DATA AND REAGENT AVAILABILITY

Strains and plasmids are available upon request. The authors affirm that all data necessary for confirming the conclusions of the article are present within the article, figures, and tables.

## ACKNOWLEDGMENTS

We thank Joshua Paulson for help in preparation of microbial growth media. We thank Laura Vanderploeg for help with figures. We thank Max Wilkinson (Memorial Sloan Kettering Cancer Center) for helpful discussions about the structure of Cwc15 and for making figures S7A and B.

## AUTHOR CONTRIBUTIONS

NZ, DH, AAH: project conceptualization; NZ, ZF, AAH: media and reagent preparation; NZ, GB, EB, ZF, SJ, and AAH: data acquisition and analysis; DH, AAH: writing and editing of the manuscript with input from NZ.

## REFERENCES

Bai, R., R. Wan, C. Yan, Q. Jia, J. Lei et al., 2021 Mechanism of spliceosome remodeling by the ATPase/helicase Prp2 and its coactivator Spp2. Science 371 (6525).

Carrocci, T.J., and K.M. Neugebauer, 2024 Emerging and re-emerging themes in co-transcriptional pre-mRNA splicing. Mol Cell 84 (19):3656–3666.

Carrocci, T.J., J.C. Paulson, and A.A. Hoskins, 2018 Functional analysis of Hsh155/SF3b1 interactions with the U2 snRNA/branch site duplex. RNA 24 (8):1028–1040.

Carrocci, T.J., D.M. Zoerner, J.C. Paulson, and A.A. Hoskins, 2017 SF3b1 mutations associated with myelodysplastic syndromes alter the fidelity of branchsite selection in yeast. Nucleic Acids Res 45 (8):4837–4852.

Chanarat, S., M. Seizl, and K. Strasser, 2011 The Prp19 complex is a novel transcription elongation factor required for TREX occupancy at transcribed genes. Genes Dev 25 (11):1147–1158.

Chung, C.S., C.K. Tseng, H.C. Chen, and S.C. Cheng, 2025 New mechanistic insights into Prp22-mediated exon ligation and mRNA release. Nucleic Acids Res 53 (16).

Chung, C.S., H.L. Wai, C.Y. Kao, and S.C. Cheng, 2023 An ATP-independent role for Prp16 in promoting aberrant splicing. Nucleic Acids Res 51 (20):10815–10828.

Company, M., J. Arenas, and J. Abelson, 1991 Requirement of the RNA helicase-like protein PRP22 for release of messenger RNA from spliceosomes. Nature 349 (6309):487–493.

de Almeida, R.A., and R.T. O’Keefe, 2015 The NineTeen Complex (NTC) and NTC-associated proteins as targets for spliceosomal ATPase action during pre-mRNA splicing. RNA Biol 12 (2):109–114.

Eysmont, K., K. Matylla-Kulinska, A. Jaskulska, M. Magnus, and M.M. Konarska, 2019 Rearrangements within the U6 snRNA Core during the Transition between the Two Catalytic Steps of Splicing. Mol Cell 75 (3):538–548 e533.

Fica, S.M., C. Oubridge, W.P. Galej, M.E. Wilkinson, X.C. Bai et al., 2017 Structure of a spliceosome remodelled for exon ligation. Nature 542 (7641):377–380.

Fica, S.M., N. Tuttle, T. Novak, N.S. Li, J. Lu et al., 2013 RNA catalyses nuclear pre-mRNA splicing. Nature 503 (7475):229–234.

Galej, W.P., M.E. Wilkinson, S.M. Fica, C. Oubridge, A.J. Newman et al., 2016 Cryo-EM structure of the spliceosome immediately after branching. Nature 537 (7619):197–201.

Goldstein, A.L., and J.H. McCusker, 1999 Three new dominant drug resistance cassettes for gene disruption in Saccharomyces cerevisiae. Yeast 15 (14):1541–1553.

Grainger, R.J., and J.D. Beggs, 2005 Prp8 protein: at the heart of the spliceosome. RNA 11 (5):533–557.

Guthrie, C., G.R. Fink, J.N. Abelson, M.I. Simon, and ScienceDirect, 1991 Guide to yeast genetics and molecular biology. San Diego ; London: Academic Press.

Henke-Schulz, L., R. Minocha, N.H. Maier, and K. Strasser, 2024 The Prp19C/NTC subunit Syf2 and the Prp19C/NTC-associated protein Cwc15 function in TREX occupancy and transcription elongation. RNA 30 (7):854–865.

Hogg, R., J.C. McGrail, and R.T. O’Keefe, 2010 The function of the NineTeen Complex (NTC) in regulating spliceosome conformations and fidelity during pre-mRNA splicing. Biochem Soc Trans 38 (4):1110–1115.

Hotz, H.R., and B. Schwer, 1998 Mutational analysis of the yeast DEAH-box splicing factor Prp16. Genetics 149 (2):807–815.

Hoyos-Manchado, R., F. Reyes-Martin, C. Rallis, E. Gamero-Estevez, P. Rodriguez-Gomez et al., 2017 RNA metabolism is the primary target of formamide in vivo. Sci Rep 7 (1):15895.

Kastner, B., C.L. Will, H. Stark, and R. Lührmann, 2019 Structural Insights into Nuclear pre-mRNA Splicing in Higher Eukaryotes. Cold Spring Harb Perspect Biol 11 (11).

Kaur, H., B. Groubert, J.C. Paulson, S. McMillan, and A.A. Hoskins, 2020 Impact of cancer-associated mutations in Hsh155/SF3b1 HEAT repeats 9-12 on pre-mRNA splicing in Saccharomyces cerevisiae. PLoS One 15 (4):e0229315.

Kaur, H., C. van der Feltz, Y. Sun, and A.A. Hoskins, 2022 Network theory reveals principles of spliceosome structure and dynamics. Structure 30 (1):190–200 e192.

Koodathingal, P., T. Novak, J.A. Piccirilli, and J.P. Staley, 2010 The DEAH box ATPases Prp16 and Prp43 cooperate to proofread 5’ splice site cleavage during pre-mRNA splicing. Mol Cell 39 (3):385–395.

Lardelli, R.M., J.X. Thompson, J.R. Yates, 3rd, and S.W. Stevens, 2010 Release of SF3 from the intron branchpoint activates the first step of pre-mRNA splicing. RNA 16 (3):516–528.

Lesser, C.F., and C. Guthrie, 1993 Mutational analysis of pre-mRNA splicing in Saccharomyces cerevisiae using a sensitive new reporter gene, CUP1. Genetics 133 (4):851–863.

Li, G.W., M.E. Wilkinson, and D.P. Bartel, 2026 The spliceosome assembles on excised linear introns to protect them from degradation. bioRxiv:2026.2001.2021.700889.

Li, Y., P. Fischer, M. Wang, Q. Zhou, A. Song et al., 2025 Structural insights into spliceosome fidelity: DHX35-GPATCH1-mediated rejection of aberrant splicing substrates. Cell Res 35 (4):296–308.

Lipinski, K.A., K.A. Senn, N.J. Zeps, and A.A. Hoskins, 2023 Biochemical and genetic evidence supports Fyv6 as a second-step splicing factor in Saccharomyces cerevisiae. RNA 29 (11):1792–1802.

Liu, L., C.C. Query, and M.M. Konarska, 2007 Opposing classes of prp8 alleles modulate the transition between the catalytic steps of pre-mRNA splicing. Nat Struct Mol Biol 14 (6):519–526.

Lleres, D., M. Denegri, M. Biggiogera, P. Ajuh, and A.I. Lamond, 2010 Direct interaction between hnRNP-M and CDC5L/PLRG1 proteins affects alternative splice site choice. EMBO Rep 11 (6):445–451.

Lu, X., and R.J. Legerski, 2007 The Prp19/Pso4 core complex undergoes ubiquitylation and structural alterations in response to DNA damage. Biochem Biophys Res Commun 354 (4):968–974.

Mayas, R.M., H. Maita, and J.P. Staley, 2006 Exon ligation is proofread by the DExD/H-box ATPase Prp22p. Nat Struct Mol Biol 13 (6):482–490.

McPheeters, D.S., 1996 Interactions of the yeast U6 RNA with the pre-mRNA branch site. RNA 2 (11):1110–1123.

Meng, E.C., T.D. Goddard, E.F. Pettersen, G.S. Couch, Z.J. Pearson et al., 2023 UCSF ChimeraX: Tools for structure building and analysis. Protein Sci 32 (11):e4792.

Ohi, M.D., A.J. Link, L. Ren, J.L. Jennings, W.H. McDonald et al., 2002 Proteomics analysis reveals stable multiprotein complexes in both fission and budding yeasts containing Myb-related Cdc5p/Cef1p, novel pre-mRNA splicing factors, and snRNAs. Mol Cell Biol 22 (7):2011–2024.

Plaschka, C., A.J. Newman, and K. Nagai, 2019 Structural Basis of Nuclear pre-mRNA Splicing: Lessons from Yeast. Cold Spring Harb Perspect Biol 11 (5).

Query, C.C., and M.M. Konarska, 2004 Suppression of multiple substrate mutations by spliceosomal prp8 alleles suggests functional correlations with ribosomal ambiguity mutants. Mol Cell 14 (3):343–354.

Query, C.C., and M.M. Konarska, 2012 CEF1/CDC5 alleles modulate transitions between catalytic conformations of the spliceosome. RNA 18 (5):1001–1013.

Rauhut, R., P. Fabrizio, O. Dybkov, K. Hartmuth, V. Pena et al., 2016 Molecular architecture of the Saccharomyces cerevisiae activated spliceosome. Science 353 (6306):1399–1405.

Sales-Lee, J., D.S. Perry, B.A. Bowser, J.K. Diedrich, B. Rao et al., 2021 Coupling of spliceosome complexity to intron diversity. Curr Biol 31 (22):4898–4910 e4894.

Schmitzova, J., C. Cretu, C. Dienemann, H. Urlaub, and V. Pena, 2023 Structural basis of catalytic activation in human splicing. Nature 617 (7962):842–850.

Schwer, B., and T. Meszaros, 2000 RNA helicase dynamics in pre-mRNA splicing. EMBO J 19 (23):6582–6591.

Senn, K.A., K.A. Lipinski, N.J. Zeps, A.F. Griffin, M.E. Wilkinson et al., 2024 Control of 3’ splice site selection by the yeast splicing factor Fyv6. Elife 13.

Silva, D., G. Santos, M. Barroca, D. Costa, and T. Collins, 2023 Inverse PCR for Site-Directed Mutagenesis. Methods Mol Biol 2967:223–238.

Slane, D., C.H. Lee, M. Kolb, C. Dent, Y. Miao et al., 2020 The integral spliceosomal component CWC15 is required for development in Arabidopsis. Sci Rep 10 (1):13336.

Soni, K., A. Horvath, O. Dybkov, M. Schwan, S. Trakansuebkul et al., 2025 Structures of aberrant spliceosome intermediates on their way to disassembly. Nat Struct Mol Biol 32 (5):914–925.

Sonstegard, T.S., J.B. Cole, P.M. VanRaden, C.P. Van Tassell, D.J. Null et al., 2013 Identification of a nonsense mutation in CWC15 associated with decreased reproductive efficiency in Jersey cattle. PLoS One 8 (1):e54872.

Sun, B., J. Huang, L. Kong, C. Gao, F. Zhao et al., 2024 Alternative splicing of a potato disease resistance gene maintains homeostasis between growth and immunity. Plant Cell 36 (9):3729–3750.

Tseng, C.K., H.L. Liu, and S.C. Cheng, 2011 DEAH-box ATPase Prp16 has dual roles in remodeling of the spliceosome in catalytic steps. RNA 17 (1):145–154.

van der Feltz, C., B. Nikolai, C. Schneider, J.C. Paulson, X. Fu et al., 2021 Saccharomyces cerevisiae Ecm2 modulates the catalytic steps of pre-mRNA splicing. RNA 27 (5):591–603.

Wan, R., R. Bai, C. Yan, J. Lei, and Y. Shi, 2019 Structures of the Catalytically Activated Yeast Spliceosome Reveal the Mechanism of Branching. Cell 177 (2):339–351 e313.

Wan, R., C. Yan, R. Bai, J. Lei, and Y. Shi, 2017 Structure of an Intron Lariat Spliceosome from Saccharomyces cerevisiae. Cell 171 (1):120–132 e112.

Wilkinson, M.E., S.M. Fica, W.P. Galej, and K. Nagai, 2021 Structural basis for conformational equilibrium of the catalytic spliceosome. Mol Cell 81 (7):1439–1452 e1439.

Wlodaver, A.M., and J.P. Staley, 2014 The DExD/H-box ATPase Prp2p destabilizes and proofreads the catalytic RNA core of the spliceosome. RNA 20 (3):282–294.

Yan, C., R. Wan, and Y. Shi, 2019 Molecular Mechanisms of pre-mRNA Splicing through Structural Biology of the Spliceosome. Cold Spring Harb Perspect Biol 11 (1).

