## Supplemental Data for "*S. cerevisiae* Cwc15p Tunes the Spliceosome Active Site for Remodeling and Catalysis"

**Supplemental Figures S1 – S7**

**Supplemental Tables S1 and S2**

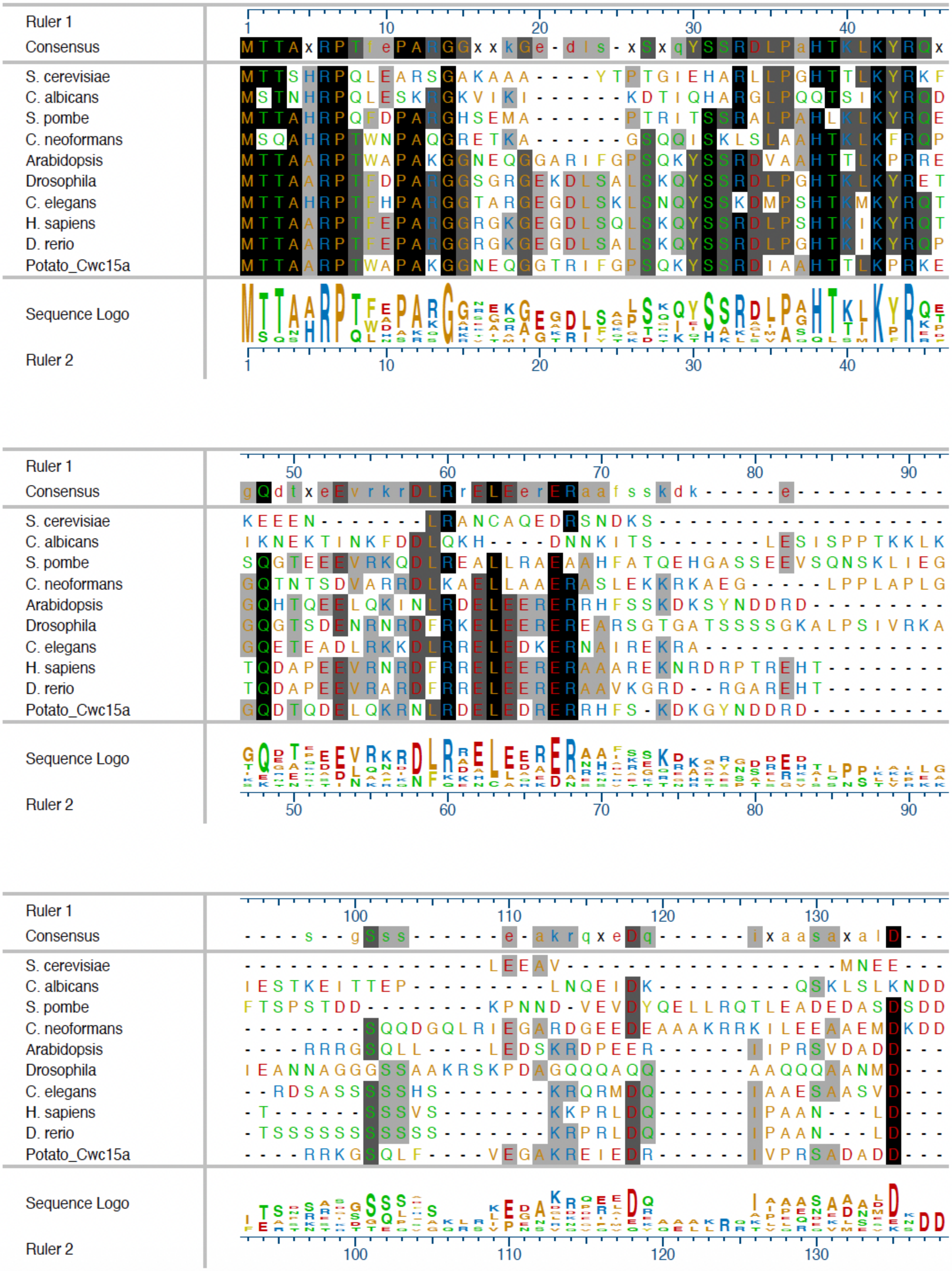

Ruler 1
Consensus
MTTAxRPTfePARGGxxkGe-dls-xSxqYSSRDLPaHTKLKYRQx
S. cerevisiae MTTSHRPQLEARSGAKAAA----YTPTGIEHARLLPGHTTLKYRKF
C. albicans MSTNHRPQLESKRGKVIKI------KDTIQHARGLPQQTSIKYRQD
S. pombe MTTAHRPQFDPARGHSEMA------PTRITSSRALPAHLKLKYRQE
C. neoformans MSQAHRPTWNPAQGRETKA------GSQQISKLSLAAHTKLKFRQP
Arabidopsis MTTAARPTWAPAKGGNEQGGARIFGPSQKYSSRDVAAHTTLKPRRE
Drosophila MTTAARPTFDPARGGSGRGEKDLSALSKQYSSRDLPGHTKLKYRET
C. elegans MTTAHRPTFHPARGGTARGEGDLSKLSNQYSSKDMPSHTKMKYRQT
H. sapiens MTTAARPTFEPARGGRGKGEGDLSQLSKQYSSRDLPSHTKIKYRQT
D. rerio MTTAARPTFEPARGGRGKGEGDLSALSKQYSSRDLPGHTKIKYRQP
Potato_Cwc15a MTTAARPTWAPAKGGNEQGGTRIFGPSQKYSSRDIAAHTTLKPRKE
Sequence Logo
Ruler 2
Ruler 1
Consensus
gQdtxeEvrkrDLRrELEerERaafssskdk-----e-----------
S. cerevisiae KEEEN-------LRANCAQEDRSNDKS-------------------
C. albicans IKNEKTINKFDDLQKH----DNNKITS-------LESISPPTKKLK
S. pombe SQGTEEEVRKQDLREALLRAEAAHFATQEHGASSEEVSQNSKLIEG
C. neoformans GQTNTSDVARRDLKAELLAAERASLEKKRKAEG-----LPPLAPLG
Arabidopsis GQHTQEELQKINLRDELEERERRHFSSKDKSYNDDRD---------
Drosophila GQGTSDENRNRDFRKELEERERERARSGTGATSSSSGKALPSIVRKA
C. elegans GQETEADLRKKDLRRELEDKERNAIREKRA----------------
H. sapiens TQDAPEEVRNRDFRRELEERERAAAREKNRDRPTREHT--------
D. rerio TQDAPEEVRARDFRRELEERERAAVKGRD--RGAREHT--------
Potato_Cwc15a GQDTQDELQKRNLRDELEDRERRHFS-KDKGYNDDRD---------
Sequence Logo
Ruler 2
Ruler 1
Consensus
-----s--gSss------e-akrqxeDq------ixaasaxalD---
S. cerevisiae ------------------LEEAV-------------------MNEE---
C. albicans IESTKEITTEP---------LNQEIDK----------QSKLSLKNDD
S. pombe FTSPSTDD--------KPNND-VEVDYQELLRQTLEADEDASDSDD
C. neoformans --------SQQDGQLRIEGARDGEEDEAAAKRRKILEEAAEMDKDD
Arabidopsis ----RRRGSQLL----LEDSKRDPEER------IIPRSVDADD---
Drosophila IEANNAGGGSSAAKRSKPDAGQQQAQQ------AAQQQAANMD---
C. elegans --RDSASSSSSHS-------KRQRMDQ------IAAESAASVD---
H. sapiens -T------SSSVS-------KKPRLDQ------IPAAN---LD---
D. rerio -TSSSSSSSSSSS-------KRPRLDQ------IPAAN---LD---
Potato_Cwc15a ----RRKGSQLF----VEGAKREIEDR------IVPRSADADD---
Sequence Logo
Ruler 2

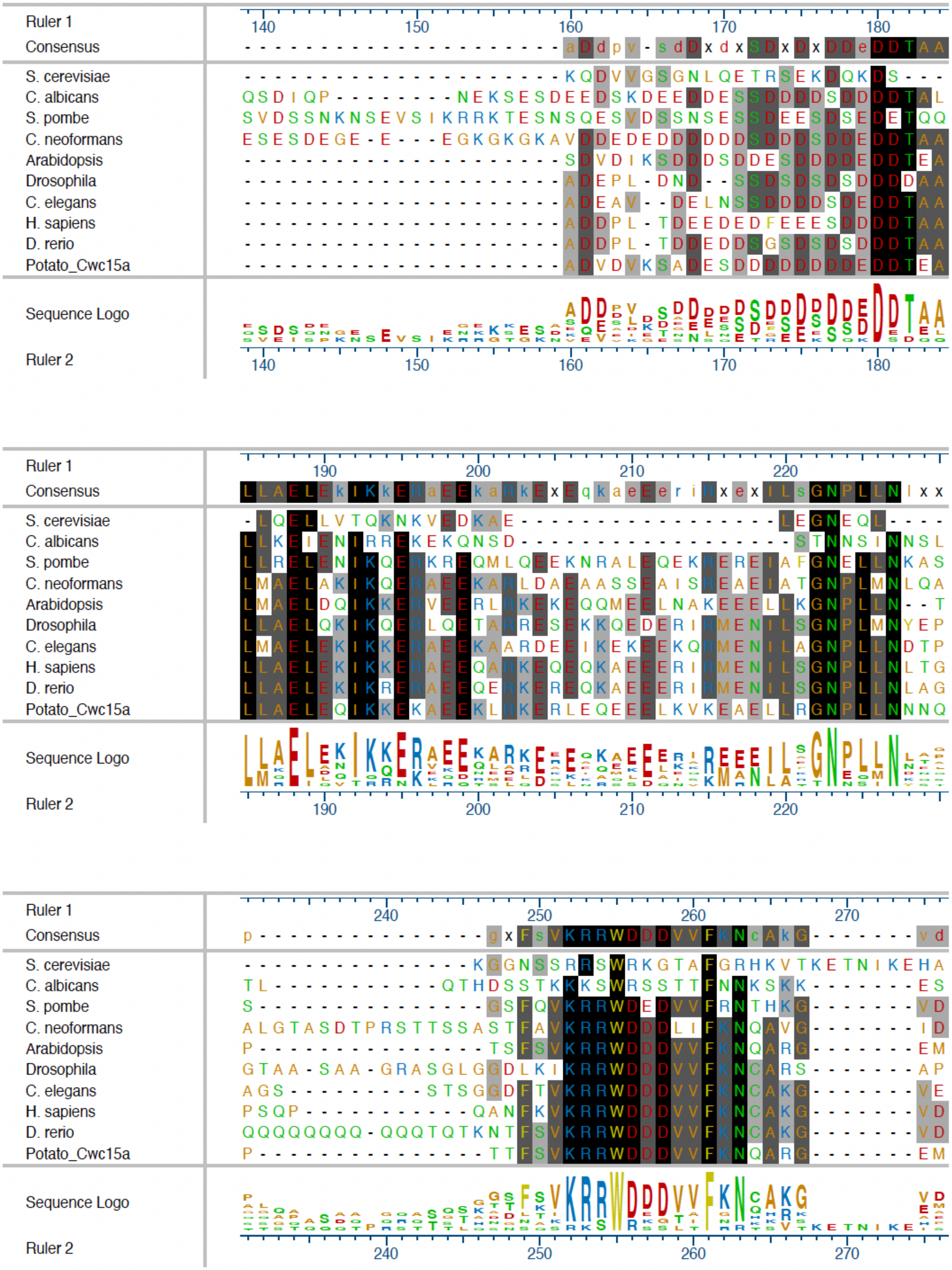
Ruler 1
Consensus
S. cerevisiae
C. albicans
S. pombe
C. neoformans
Arabidopsis
Drosophila
C. elegans
H. sapiens
D. rerio
Potato_Cwc15a
Sequence Logo
Ruler 2
140
150
160
170
180
190
200
210
220
240
250
260
270

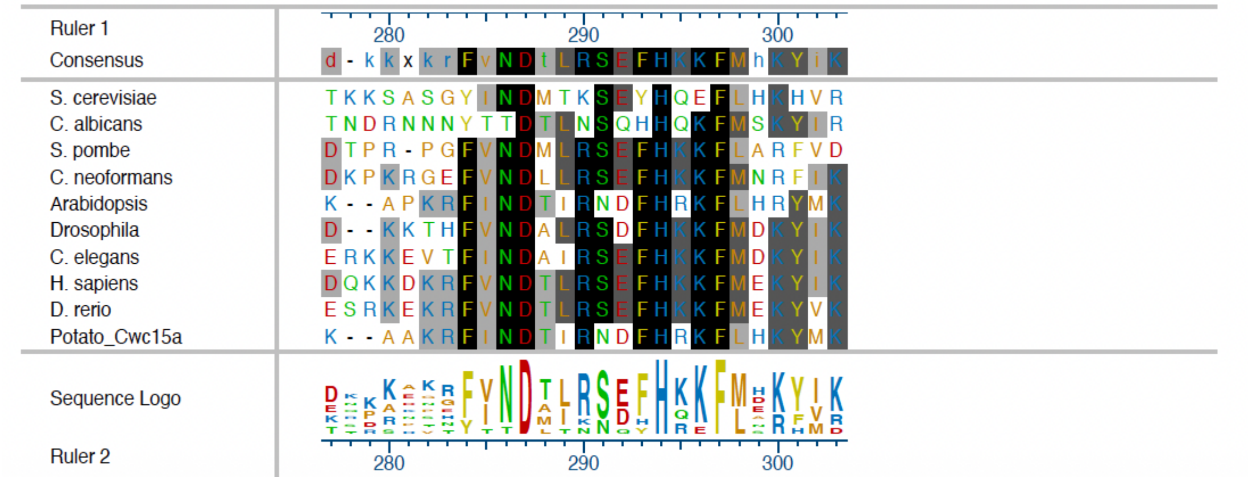

**Supplemental Figure S1.** Sequence alignment of Cwc15 proteins. The alignment was prepared using Clustal Omega implemented in MegAlign Pro (Lasergene, DNASTAR, Inc.).

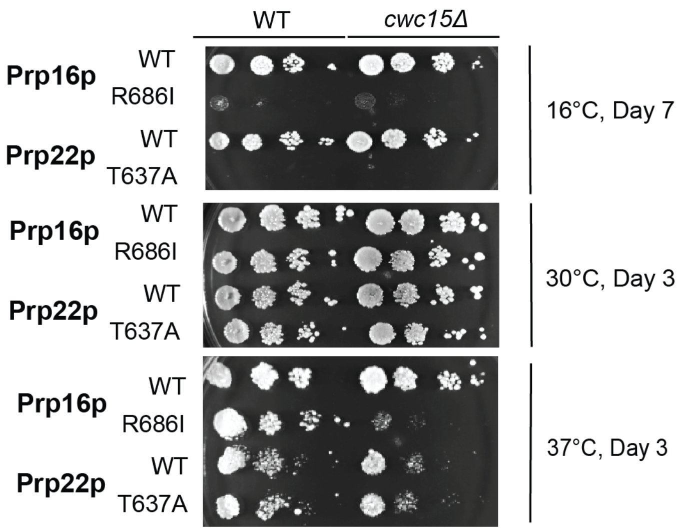

**Supplemental Figure S2.** Additional replicate of plating assays testing genetic interactions between *cwc15*Δ and Prp16 or Prp22 mutants from **Fig. 2C**.

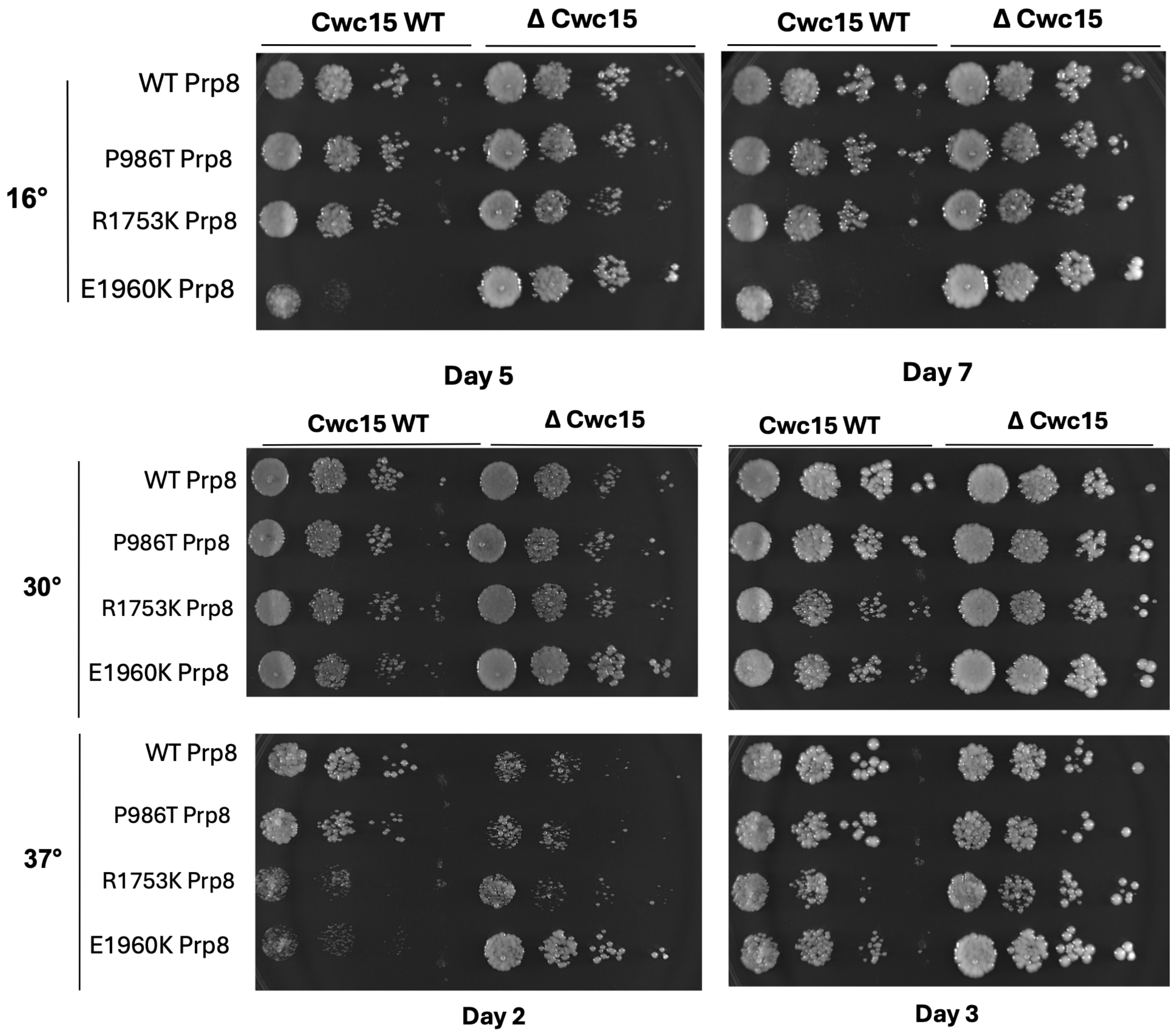

**Supplemental Figure S3** Additional replicate of plating assays testing genetic interactions between *cwc15*Δ and select Prp8 mutants from **Fig. 3B** at three different temperatures. Note increased growth for *cwc15*Δ yeast in combination with R1753K or E1960K mutations of Prp8 relative to *CWC15* (WT) yeast at 16 and 37$^{o}$C.

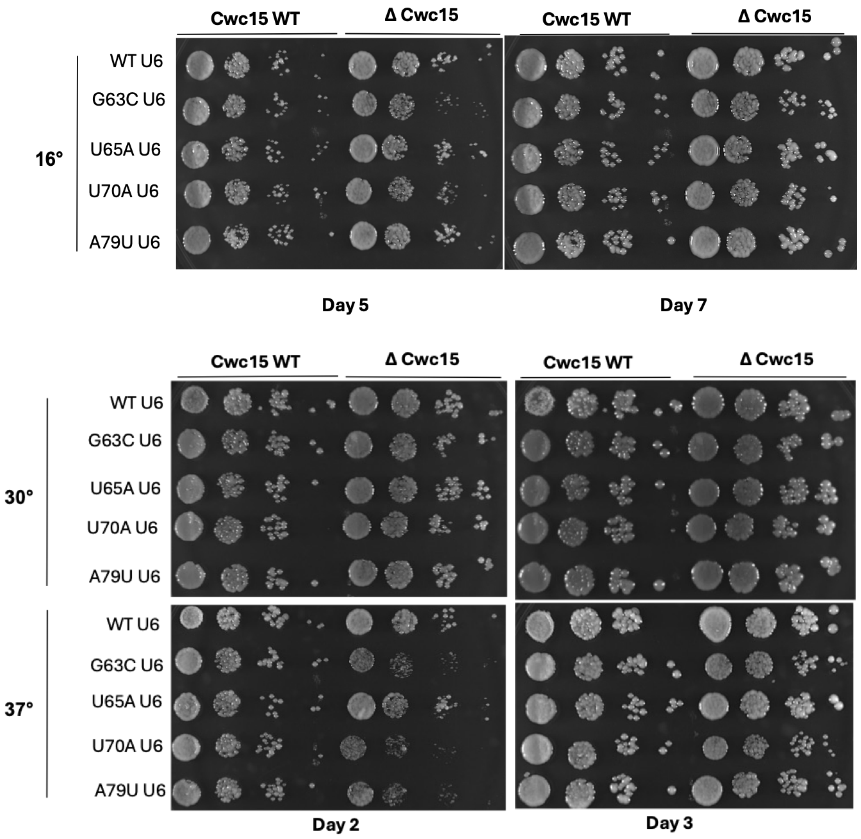

**Supplemental Figure S4.** Additional replicate of plating assays testing genetic interactions between *cwc15*Δ and select U6 mutants from **Fig. 3C** at three temperatures. Note that *cwc15*Δ yeast grow more slowly at 37°C in combination with U6-C63C, U70A, and A79U mutations.

A

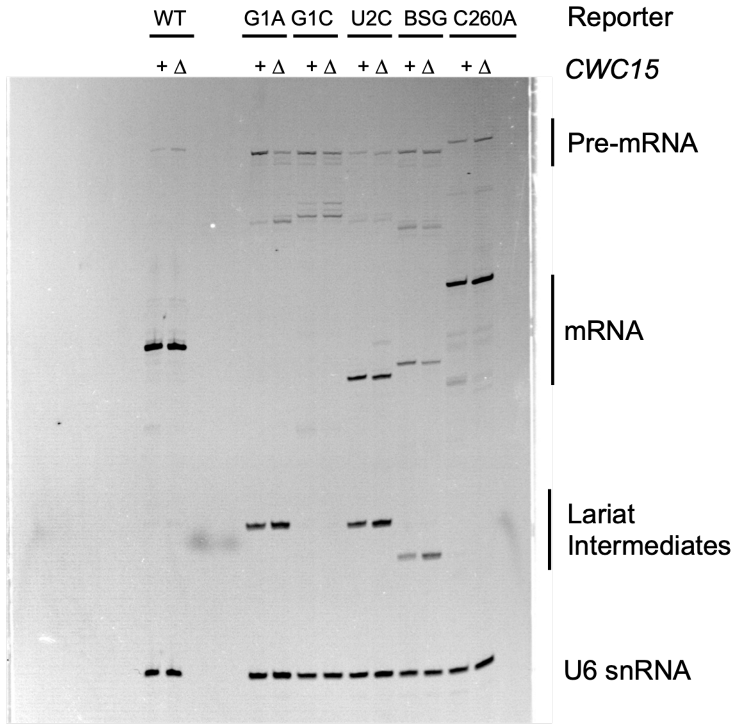

B

| Reporter | Strain | Ratio of mRNA/U6 |
|---|---|---|
| WT | WT | 1 |
| | *cwc15Δ* | 0.93 |
| G1A | WT | ND |
| | *cwc15Δ* | ND |
| G1C | WT | ND |
| | *cwc15Δ* | ND |
| U2C | WT | 0.26 |
| | *cwc15Δ* | 0.20 |
| BSG | WT | 0.14 |
| | *cwc15Δ* | 0.08 |
| C260A | WT | 0.57 |
| | *cwc15Δ* | 0.60 |

C

| Reporter | Strain | Ratio of mRNA/total |
|---|---|---|
| WT | WT | 1 |
| | *cwc15Δ* | 0.99 |
| G1A | WT | ND |
| | *cwc15Δ* | ND |
| G1C | WT | ND |
| | *cwc15Δ* | ND |
| U2C | WT | 1 |
| | *cwc15Δ* | 0.77 |
| BSG | WT | 1 |
| | *cwc15Δ* | 0.47 |
| C260A | WT | 1 |
| | *cwc15Δ* | 0.99 |

**Supplemental Figure S5.** Primer extension analysis of ACT1-CUP1 RNAs isolated from WT or *cwc15*Δ yeast. The U6 snRNA was used as a control. Note that the reporters encoded RNAs of different sizes, and that the G1A and G1C mRNAs are too low in abundance to have been visualized. For (B), the integrated intensities of each mRNA band was compared to that of each U6 band and then normalized to results obtained with the WT ACT1-CUP1 reporter and the WT yeast stain. For (C), the ratio of mRNA/(pre-mRNA+mRNA+Lariat Intermediate) was determined and results from each *cwc15Δ* strain were normalized to the corresponding WT strain.

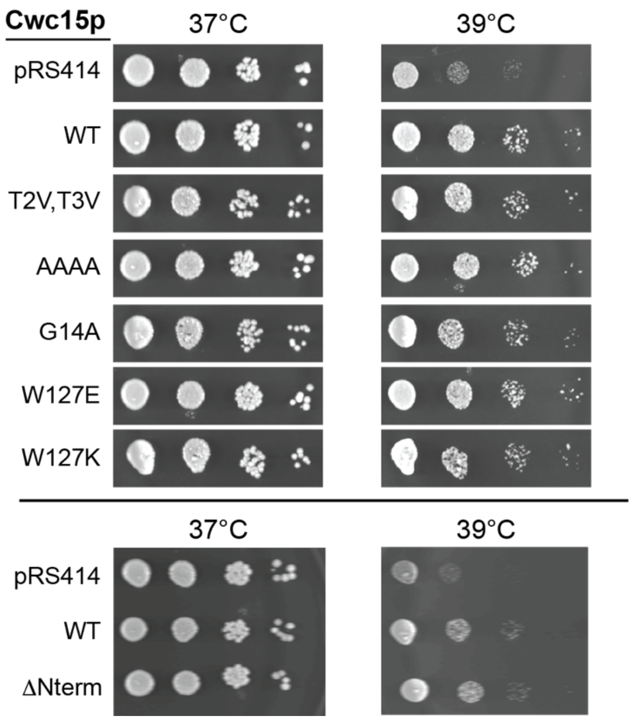

**Supplemental Figure S6.** Growth assays of *cwc15*Δ yeast expressing Cwc15p mutants relative to WT and empty plasmid (pRS414) controls. Yeast were grown on -TRP DO plates and imaged after 2 days.

.

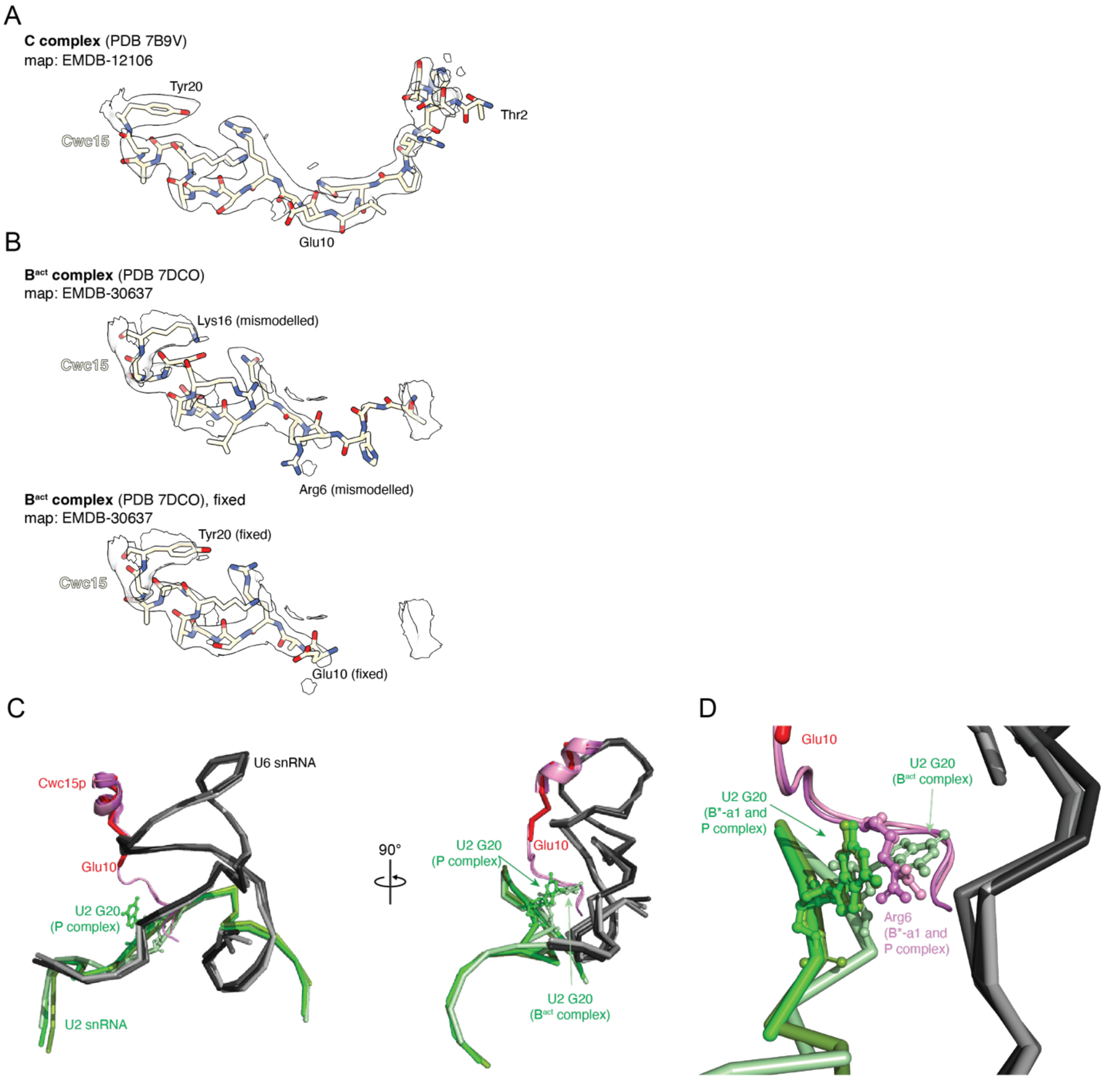

**Figure S7. Alignment of Cwc15p N-terminus Structures from Cryo-EM Models of Spliceosomes.** (**A**) Superposition of the EM density and model of the Cwc15p N-terminus from the of the C complex spliceosome at 2.8 Å. (**B**) (Top) Superposition of the EM density and PDB-deposited model for the Cwc15p N-terminus from the $B^{act}$ complex spliceosome at 2.5 Å. Note the similarities in density shapes with those shown in panel (A) especially around Lys16 (similar to Tyr20 in panel A) and Gln8 (similar to Arg12 in panel A). (Bottom) A corrected model for the N-terminus of Cwc15p in the $B^{act}$ spliceosome. The N-terminal nine amino acids could not be modeled due to a lack of specific density. (**C**) Superposition of active sites from $B^{act}$ (7DCO, corrected), $B^{*\text{-a1}}$ (6J6H), and P complex (9DTR) spliceosomes. The U6 snRNAs are shown in grey/black colors, the U2 snRNA in green colors, and Cwc15p proteins red/pink colors. Cwc15p Glu10 (the last modeled residue in the corrected $B^{act}$ structure) is labeled. Note that the

U2 snRNA backbone changes conformation relative to U6 between $B^{act}$ and later structures and that U2 G20 moves from being positioned in the cleft between U2 and U6 to being located outside the cleft. (**D**) When U2 G20 is located in the cleft ($B^{act}$), it may sterically occlude insertion of the Cwc15p N-terminus. However, when G20 is outside the cleft ($B^{*-a1}$ and P) it can form a cation-$\pi$ interaction with Cwc15p Arg6. Figure panels A-C were made using ChimeraX (Meng et al. 2023). Figure panels C and D were made using Pymol (Schrödinger).

### **Supplemental Table S1.** Yeast strains

| Lab ID | Genotype | Plasmid | Notes | Figures | Notes/Ref. |
|---|---|---|---|---|---|
| yAAH0396 | MATa his3 leu2Δ0 met15Δ 0 ura3Δ0 | -- | BY4741 | 1C | |
| yAAH3574 | yAAH0396 + *cwc15Δ*::HygR | -- | *cwc15Δ* | 1C | |
| yAAH3767 | yAAH0135 + (PRP2/CEN/LEU2) + *cwc15Δ*::HygR | pAAH0778 | Prp2 Shuffle | 2B | |
| yAAH3768 | yAAH0135 + (PRP2-Q548N/CEN/LEU2) + *cwc15Δ*::HygR | pAAH0790 | Prp2 Shuffle | 2B | |
| yAAH0135 | PRP2::TRP1 lys2-801 his3- Δ 200 leu2-3,112 trp1-1 ura3-52 (YCp50-Prp2) | pAAH0770 | Prp2 Shuffle | 2B | Gift of R.J. Lin |
| yAAH0130 | MATa trp1 ura3 his3 lys2 leu2 ade2 prp16Δ::lys2 pSB-Prp16 | pAAH0069 | Prp16 Shuffle | 2B | Gift of C. Guthrie (aka yS78) |
| yAAH3770 | yAAH0130 + (PRP16-R686I/CEN/TRP1) + *cwc15Δ*::HygR | pAAH1039 | Prp16 Shuffle | 2B | |
| yAAH3769 | yAAH0130 + (PRP16/CEN/TRP1) + *cwc15Δ*::HygR | pAAH1040 | Prp16 Shuffle | 2B | |
| yAAH3824 | yAAH1930 + (PRP22-T637A/CEN/TRP1) + *cwc15Δ*::HygR | pAAH1043 | Prp22 Shuffle | 2B | |
| yAAH3823 | yAAH1930 + (PRP22 /CEN/TRP1) + *cwc15Δ*::HygR | pAAH1042 | Prp22 Shuffle | 2B | |
| yAAH1930 | MATa ade2 cup1Δ::ura3 his3 leu2 lys2 trp1 ura3 GAL prp22Δ:loxP + (PRP22/CEN/URA3) | PRP22/URA | Prp22 Shuffle | 2B | Gift of Charles Query and Magda Konarska (aka yMK-02) |
| pAAH3809 | yAAH0117 + (PRP8/TRP1/2μ) + *cwc15Δ*::HygR | pAAH0997 | Prp8 Shuffle | 3B | |
| yAAH3810 | yAAH0117 + (PRP8-P986T/TRP1/2μ) + *cwc15Δ*::HygR | pAAH1001 | Prp8 Shuffle | 3B | |
| yAAH3812 | yAAH0117 + (PRP8-R1753K/TRP1/2μ) + *cwc15Δ*::HygR | pAAH1004 | Prp8 Shuffle | 3B | |
| yAAH3813 | yAAH0117 + (PRP8-P986T+R1753K/TRP1/2μ) + *cwc15Δ*::HygR | pAAH1006 | Prp8 Shuffle | 3B | |
| yAAH3811 | yAAH0117 + (PRP8-E1960K/TRP1/2μ) + *cwc15Δ*::HygR | pAAH1003 | Prp8 Shuffle | 3B | |
| yAAH0117 | ade2 cup1Δ::ura3 his 3 leu2 lys2 prp8 Δ::lys2 trp1 pJU169 (PRP8/URA3) | pAAH0088 | Prp8 Shuffle | 3B | Gift of C. Guthrie (aka yJU75) |
| yAAH3826 | yAAH1908 + (SNR6-U70A/TRP1/CEN) + *cwc15Δ*::HygR | pAAAH1118 | U6 Shuffle | 3C | |
| yAAH3827 | yAAH1908 + (SNR6-G63C/TRP1/CEN) + *cwc15Δ*::HygR | pAAH1119 | U6 Shuffle | 3C | |
| yAAH3828 | yAAH1908 + (SNR6-A79U/TRP1/CEN) + *cwc15Δ*::HygR | pAAH1120 | U6 Shuffle | 3C | |
| yAAH3840 | yAAH1908 + (SNR6/TRP1/CEN) + *cwc15Δ*::HygR | pAAH0412 | U6 Shuffle | 3C | |
| yAAH3841 | yAAH1908 + (SNR6-U57C/TRP1/CEN) + *cwc15Δ*::HygR | pAAH1028 | U6 Shuffle | 3C | |
| yAAH3842 | yAAH1908 + (SNR6-U57A/TRP1/CEN) + *cwc15Δ*::HygR | pAAH1027 | U6 Shuffle | 3C | |
| yAAH3849 | yAAH1908 + (SNR6-U65A/TRP1/CEN) + *cwc15Δ*::HygR | pAAH1686 | U6 Shuffle | 3C | |
| yAAH1908 | MATα cup1Δ ura3 his3 trp1 lys2 ade2 leu2 snr6::NatR (SNR6 SNR4/URA3/CEN) | pAAH0989 | U6 Shuffle | 3C | |
| yAAH3575 | yAAH0434 + *cwc15Δ*::HygR | -- | CUP1 | 4B | |
| yAAH0434 | MATα cup1Δ ura3 his3 trp1 lys2 ade2 leu2 | -- | CUP1 | 4B | Gift of D. Brow (aka 46α) |
| yAAH2852 | BY4741 (MATa MATa his3 leu2Δ0 met15Δ 0 ura3Δ0) + trp1Δ63 | -- | -- | S6 | BY4741 trp1 mutant |
| yAAH4041 | yAAH2852 + *cwc15Δ*::HygR | -- | *cwc15Δ* | S6 | |
| yAAH4042 | + pRS414 (TRP1/CEN) | pAAH0135 | *cwc15Δ* | S6 | |
| yAAH4043 | + (WT CWC15/TRP1/CEN) | pAAH1687 | *cwc15Δ* | S6 | |
| yAAH4044 | + (T2V T3V CWC15/TRP1/CEN) | pAAH1756 | *cwc15Δ* | S6 | |
| yAAH4045 | + (AAAA CWC15/TRP1/CEN) | pAAH1746 | *cwc15Δ* | S6 | |

| | | | | |
|---|---|---|---|---|
| yAAH4046 | + (G14A CWC15/TRP1/CEN) | pAAH1747 | *cwc15Δ* | S6 |
| yAAH4047 | + (W127E CWC15/TRP1/CEN) | pAAH1749 | *cwc15Δ* | S6 |
| yAAH4048 | + (W127K CWC15/TRP1/CEN) | pAAH1748 | *cwc15Δ* | S6 |
| yAAH4078 | +(NtermΔ CWC15/TRP1/CEN) | pAAH1748 | *cwc15Δ* | S6 |
| yAAH3844 | yAAH1908 + (SNR6-WT/TRP1/CEN) + *cwc15Δ*::HygR | pAAH0412 | U6 Shuffle | 3C,6B |
| yAAH3919 | yAAH1908 + (SNR6-A62U/TRP1/CEN) | pAAH1738 | U6 Shuffle | 6B |
| yAAH3902 | yAAH1908 + (SNR6-A62U/TRP1/CEN) + *cwc15Δ*::HygR | pAAH1738 | U6 Shuffle | 6B |
| yAAH3920 | yAAH1908 + (SNR6-G63U/TRP1/CEN) | pAAH1739 | U6 Shuffle | 6B |
| yAAH3903 | yAAH1908 + (SNR6-G63U/TRP1/CEN) + *cwc15Δ*::HygR | pAAH1739 | U6 Shuffle | 6B |
| yAAH3921 | yAAH1908 + (SNR6-C84G/TRP1/CEN) | pAAH1740 | U6 Shuffle | 6B |
| yAAH3904 | yAAH1908 + (SNR6-C84G/TRP1/CEN) + *cwc15Δ*::HygR | pAAH1740 | U6 Shuffle | 6B |
| yAAH3922 | yAAH1908 + (SNR6-C85G/TRP1/CEN) | pAAH1741 | U6 Shuffle | 6B |
| yAAH3905 | yAAH1908 + (SNR6-C85G/TRP1/CEN) + *cwc15Δ*::HygR | pAAH1741 | U6 Shuffle | 6B |
| yAAH4079 | yAAH0117 + (PRP8-P986T+R1753K/TRP1/2μ) + *cwc15Δ*::HygR + pRS413 (HIS3/CEN) | pRS413 | *cwc15Δ*/ Prp8 Shuffle | 5E |
| yAAH4080 | yAAH0117 + (PRP8-P986T+R1753K/TRP1/2μ) + *cwc15Δ*::HygR (WT CWC15/ HIS3/CEN) | pAAH1865 | *cwc15Δ*/ Prp8 Shuffle | 5E |
| yAAH4081 | yAAH0117 + (PRP8-P986T+R1753K/TRP1/2μ) + *cwc15Δ*::HygR + (T2V T3V CWC15/ HIS3/CEN) | pAAH1866 | *cwc15Δ*/ Prp8 Shuffle | 5E |
| yAAH4082 | yAAH0117 + (PRP8-P986T+R1753K/TRP1/2μ) + *cwc15Δ*::HygR + (AAAA CWC15/ HIS3/CEN) | pAAH1867 | *cwc15Δ*/ Prp8 Shuffle | 5E |
| yAAH4083 | yAAH0117 + (PRP8-P986T+R1753K/TRP1/2μ) + *cwc15Δ*::HygR + (G14A CWC15/ HIS3/CEN) | pAAH1868 | *cwc15Δ*/ Prp8 Shuffle | 5E |
| yAAH4084 | yAAH0117 + (PRP8-P986T+R1753K/TRP1/2μ) + *cwc15Δ*::HygR + (W127E CWC15/ HIS3/CEN) | pAAH1869 | *cwc15Δ*/ Prp8 Shuffle | 5E |
| yAAH4085 | yAAH0117 + (PRP8-P986T+R1753K/TRP1/2μ) + *cwc15Δ*::HygR + (W127K CWC15/ HIS3/CEN) | pAAH1870 | *cwc15Δ*/ Prp8 Shuffle | 5E |
| yAAH4086 | yAAH0117 + (PRP8-P986T+R1753K/TRP1/2μ) + *cwc15Δ*::HygR +(NtermΔ CWC15/ HIS3/CEN) | pAAH1871 | *cwc15Δ*/ Prp8 Shuffle | 5E |
| yAAH4087 | yAAH1908 + (SNR6-U57C/TRP1/CEN) + *cwc15Δ*::HygR + pRS413 (HIS3/CEN) | pRS413 | *cwc15Δ*/ U6 Shuffle | 5F |
| yAAH4088 | yAAH1908 + (SNR6-U57C/TRP1/CEN) + *cwc15Δ*::HygR + (WT CWC15/HIS3/CEN) | pAAH1865 | *cwc15Δ*/ U6 Shuffle | 5F |
| yAAH4089 | yAAH1908 + (SNR6-U57C/TRP1/CEN) + *cwc15Δ*::HygR + (T2V T3V CWC15/HIS3/CEN) | pAAH1866 | *cwc15Δ*/ U6 Shuffle | 5F |
| yAAH4090 | yAAH1908 + (SNR6-U57C/TRP1/CEN) + *cwc15Δ*::HygR + (AAAA CWC15/HIS3/CEN) | pAAH1867 | *cwc15Δ*/ U6 Shuffle | 5F |
| yAAH4091 | yAAH1908 + (SNR6-U57C/TRP1/CEN) + *cwc15Δ*::HygR + (G14A CWC15/HIS3/CEN) | pAAH1868 | *cwc15Δ*/ U6 Shuffle | 5F |

| | | | | |
|---|---|---|---|---|
| yAAH4092 | yAAH1908 + (SNR6-U57C/TRP1/CEN) + *cwc15Δ*::HygR + (W127E CWC15/HIS3/CEN) | pAAH1869 | *cwc15Δ*/ U6 Shuffle | 5F |
| yAAH4093 | yAAH1908 + (SNR6-U57C/TRP1/CEN) + *cwc15Δ*::HygR + (W127K CWC15/HIS3/CEN) | pAAH1870 | *cwc15Δ*/ U6 Shuffle | 5F |
| yAAH4094 | yAAH1908 + (SNR6-U57C/TRP1/CEN) + *cwc15Δ*::HygR +(NtermΔ CWC15/HIS3/CEN) | pAAH1871 | *cwc15Δ*/ U6 Shuffle | 5F |

*Additional yeast strains containing CWC15 and wild-type or mutant Prp2, Prp16, Prp22, Prp8, or U6 genes corresponding to those with *cwc15*Δ have been previously reported by our group (van der Feltz et al. 2021; Senn et al. 2024; Lipinski et al. 2023).

**Supplemental Table S2.** Plasmids

| Lab ID | Description | Figures | Notes/Ref. |
|---|---|---|---|
| pAAH0778 | pRS415, WT Prp2 | 2B | Gift of D. Brow |
| pAAH0790 | pRS415, WT Prp2-Q548N | 2B | |
| pAAH0770 | yCP50, WT Prp2 | 2B | Gift of R.J. Lin |
| pAAH0069 | pSB2-16, WT Prp16 | 2B | Gift of C. Guthrie |
| pAAH1039 | Prp16-R686I/TRP/CEN | 2B | Gift of C. Query, B. Schwer |
| pAAH1040 | WT Prp16/TRP/CEN | 2B | Gift of C. Query, B. Schwer |
| pAAH1043 | Prp22-T637A/TRP/CEN | 2B | Gift of C. Query, B. Schwer |
| pAAH1042 | WT Prp22 /TRP/CEN | 2B | Gift of C. Query, B. Schwer |
| pAAH0997 | pRS424, Prp8, pJU225-4 | 3B | Gift of D. Brow |
| pAAH1001 | Prp8-161 allele, P986T/TRP, 2μ | 3B | Gift of M. Konarska |
| pAAH1004 | Prp8 R1753K/TRP, 2μ | 3B | Gift of M. Konarska |
| pAAH1006 | Prp8 P986T R1753K/TRP, 2μ | 3B | Gift of M. Konarska |
| pAAH1003 | Prp8-101 allele, E1960K/TRP, 2μ | 3B | Gift of M. Konarska |
| pAAH0088 | pJU169 WT Prp8/URA | 3B | Gift of C. Guthrie |
| pAAH1118 | U6-U70A/TRP/CEN | 3C | |
| pAAH1119 | U6-G63C/TRP/CEN | 3C | |
| pAAH1120 | U6-A79U/TRP/CEN | 3C | |
| pAAH0412 | WT U6/TRP/CEN (pRS314) | 3C, 6B | Gift of D. Brow |
| pAAH1028 | U6-U57C/TRP/CEN | 3C | |
| pAAH1027 | U6-U57A/TRP/CEN | 3C | |
| pAAH1686 | U6-U65A/TRP/CEN | 3C | |
| pAAH0989 | WT U4, WT U6/URA/CEN | 3C | Gift of D. Brow |
| pAAH0135 | pRS414 | S6 | |
| pAAH1687 | WT CWC15/TRP/CEN | S6 | Made by cloning the *S. cerevisiae* CWC15 gene (+146nt upstream and +338nt downstream) into the BamHI and EcoRI sites of pRS414 |
| pAAH1756 | pAAH1687 + T2V T3V mutants | S6 | |
| pAAH1746 | pAAH1687 + H5A, R6A, P7A, Q8A mutants | S6 | |
| pAAH1747 | pAAH1687 + G14A mutant | S6 | |
| pAAH1749 | pAAH1687 + W127E mutant | S6 | |
| pAAH1748 | pAAH1687 + W127K mutant | S6 | |
| pAAH1738 | U6-A62U/TRP/CEN | 6B | |
| pAAH1739 | U6-G63U/TRP/CEN | 6B | |
| pAAH1740 | U6-C84G/TRP/CEN | 6B | |
| pAAH1741 | U6-C85G/TRP/CEN | 6B | |
| pAAH1865 | WT CWC15/HIS3/CEN | 5 | |
| pAAH1866 | pAAH1865 + T2V T3V mutants | 5 | |
| pAAH1867 | pAAH1865 + H5A, R6A, P7A, Q8A mutants | 5 | |
| pAAH1868 | pAAH1865 + G14A mutant | 5 | |
| pAAH1869 | pAAH1865 + W127E mutant | 5 | |
| pAAH1870 | pAAH1865 + W127K mutant | 5 | |
| pAAH1871 | pAAH1865 + NtermΔ mutant | 5 | |
| pAAH1872 | pAAH1687 + NtermΔ mutant | S6 | |

### REFERENCES

Lipinski, K.A., K.A. Senn, N.J. Zeps, and A.A. Hoskins, 2023 Biochemical and genetic evidence supports Fyv6 as a second-step splicing factor in Saccharomyces cerevisiae. *RNA* 29 (11):1792–1802.

Meng, E.C., T.D. Goddard, E.F. Pettersen, G.S. Couch, Z.J. Pearson *et al.*, 2023 UCSF ChimeraX: Tools for structure building and analysis. *Protein Sci* 32 (11):e4792.

Senn, K.A., K.A. Lipinski, N.J. Zeps, A.F. Griffin, M.E. Wilkinson *et al.*, 2024 Control of 3' splice site selection by the yeast splicing factor Fyv6. *Elife* 13.

van der Feltz, C., B. Nikolai, C. Schneider, J.C. Paulson, X. Fu *et al.*, 2021 Saccharomyces cerevisiae Ecm2 modulates the catalytic steps of pre-mRNA splicing. *RNA* 27 (5):591–603.
